# RASopathy Mutations Provide Functional Insight into the BRAF Cysteine-rich Domain and Demonstrate the Importance of Autoinhibition in BRAF Regulation

**DOI:** 10.1101/2021.10.01.462773

**Authors:** Russell Spencer-Smith, Elizabeth M. Terrell, Christine Insinna, Constance Agamasu, Morgan E. Wagner, Daniel A. Ritt, Jim Stauffer, Andrew G. Stephen, Deborah K Morrison

## Abstract

BRAF is frequently mutated in human cancer and the RASopathy syndromes, with RASopathy mutations often observed in the cysteine-rich domain (CRD). Although the CRD participates in phosphatidylserine (PS) binding, the RAS-RAF interaction, and RAF autoinhibition, the impact of these activities on RAF function in normal and disease states is not well-characterized. Here, we analyze a panel of CRD mutations and show that they increase BRAF activity by relieving autoinhibition and/or enhancing PS binding, with relief of autoinhibition being the major factor determining mutation severity in zebrafish models. Further, we show that CRD-mediated autoinhibition is essential for preventing the constitutive plasma membrane localization of BRAF and increased RAS-dependent and RAS-independent function. Comparison of the BRAF- and CRAF-CRDs also indicates that the BRAF-CRD is a stronger mediator of autoinhibition and PS binding, and given the increased catalytic activity of BRAF, our studies reveal a more critical role for CRD-mediated autoinhibition in BRAF regulation.

## INTRODUCTION

RAF family kinases (ARAF, BRAF, and CRAF) are primary effectors of the RAS GTPases and essential members of the pro-proliferative/anti-apoptotic RAS-ERK MAPK pathway. The RAFs can be divided into two functional domains, the C-terminal catalytic domain (CAT) and the N-terminal regulatory (REG) domain, the latter of which contains a conserved RAS binding domain (RBD), a cysteine-rich domain (CRD), and a serine/threonine rich region (Lavoie and Therrien, 2015; Terrell and Morrison, 2019); Figure 1A). Under normal signaling conditions RAF kinases cycle between three major states: 1) an autoinhibited, monomeric conformation that localizes to the cytosol (Chong and Guan, 2003; Cutler et al., 1998; Nan et al., 2013), 2) an activated, RAS-bound dimeric configuration at the plasma membrane (Freeman et al., 2013; Hu et al., 2013); and 3) a signaling-incompetent monomeric form that has been inactivated and released from the membrane via signal-induced feedback phosphorylation (Dougherty et al., 2005; Ritt et al., 2010). This regulatory cycle can be disrupted by mutational events to promote upregulated RAF signaling in human disease states, with somatic and germline mutations being causative for cancer and the developmental RASopathy syndromes, respectively (Schubbert et al., 2007). Although the vast majority of oncogenic RAF mutations occur in *BRAF*, driver mutations in the RASopathies are observed in both *BRAF* and *CRAF* (Hebron et al., 2022).

**Figure 1.**
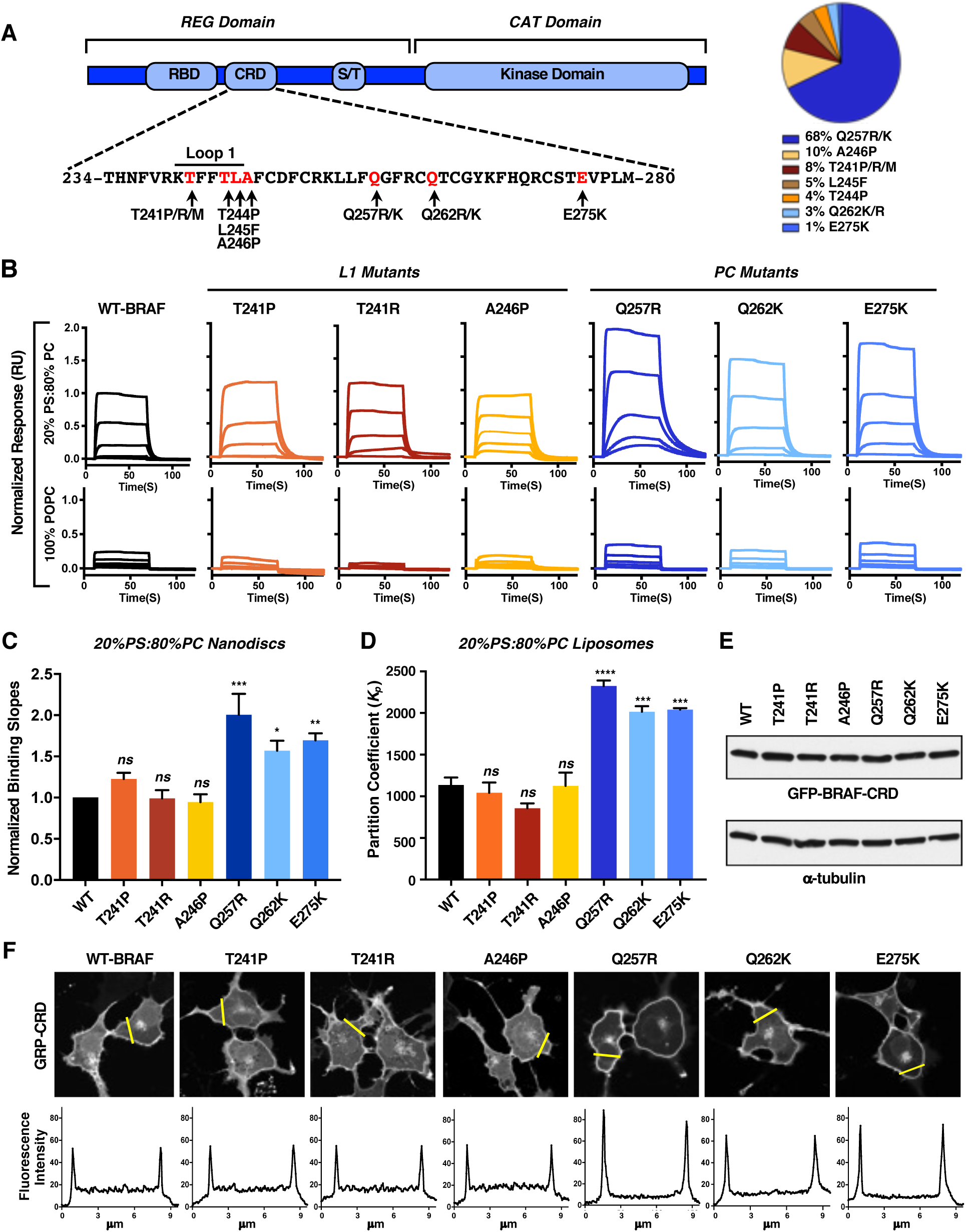
Identification of BRAF-CRD Mutations that Modulate PS Binding. (A) BRAF domain structure and the CRD amino acid sequence is shown with arrows indicating the RASopathy-associated mutations (Left panel). The percent occurrence of a particular BRAF-CRD mutation is also indicated (Right panel). Data were collected from nseuronet.com. (B) A 2-fold dilution series of purified 5 µM WT or mutant BRAF-CRD proteins was injected over nanodiscs containing 20% POPS/80% POPC or 100% POPC, and binding responses were determined by SPR. Sensorgrams were normalized to the WT-BRAF-CRD binding response. (C) Binding slopes of the CRD mutants were normalized to the binding slope of the WT-CRD (with WT equaling 1), and the graph represents the mean of 3 independent experiments ± SD. (D) Partitioning coefficients for binding of WT and mutant BRAF-CRDs to liposomes containing 20% POPS/80% POPC were determined, and the graph represents the mean of 3 independent experiments ± SD. (E, F) Serum-starved COS-7 cells transiently expressing the indicated GFP-CRD proteins were examined by immunoblot analysis (E) and by confocal microscopy (F). A tracing depicting the GFP intensity in the area indicated by the yellow line is also shown. The images are representative of 2 independent experiments with similar results. For statistical analysis (C and D), student’s *t*-test (two-tailed, assuming unequal variance) was performed. ns, not significant; *, *P* < 0.05, P **, *P* < 0.01; ***, *P* < 0.001, and ****, *P* < 0.0001.

The RASopathies are a group of related developmental disorders caused by germline mutations in components of the RAS-ERK MAPK pathway (Rauen, 2013). These disorders, which include cardiofaciocutaneous (CFC) syndrome, Costello syndrome, Noonan syndrome, and Noonan syndrome with multiple lentigines, are characterized by distinct craniofacial abnormalities, developmental delays, cardiac defects, and an increased risk of certain childhood cancers (Tidyman and Rauen, 2016). With regard to the RAF kinases, RASopathy mutations in BRAF versus CRAF tend to occur in different regions of the proteins and are primarily associated with different RASopathy syndromes. For example, CRAF RASopathy mutations are largely observed in Noonan syndrome (NS; 6% of NS cases) and typically lie in close proximity to a 14-3-3 docking site found in the RAF regulatory domain (Pandit et al., 2007; Razzaque et al., 2007). In contrast, germline alterations in BRAF are mainly associated with CFC (75% of CFC cases), and these mutations are most commonly observed either in the kinase domain (∼62% of mutations) or the cysteine-rich domain (CRD; ∼38% of mutations) (Niihori et al., 2006; Rodriguez-Viciana et al., 2006).

CRDs (also known as C1 domains) are zinc-finger domains known to mediate lipid and protein interactions (Colon-Gonzalez and Kazanietz, 2006). The ligand binding interface of a CRD consists of two flexible loops (Loop 1 and 2) that contain hydrophobic residues able to insert into the membrane bilayer and basic residues that can interact with lipid head groups (Medkova and Cho, 1999; Xu et al., 1997). Characterization of the RAF-CRDs has indicated an interaction with both phosphatidylserine and RAS, which contributes to the stable plasma membrane recruitment of RAF and is required for RAS-driven ERK activation (Bondeva et al., 2002; Brtva et al., 1995; Daub et al., 1998; Fang et al., 2020; Ghosh et al., 1996; Ghosh et al., 1994; Hekman et al., 2002; Hibino et al., 2011; Hu et al., 1995; Roy et al., 1997; Tran et al., 2021). The CRD has also been shown to play a prominent role in mediating the autoinhibitory effect of the RAF regulatory domain, which maintains RAF monomers in an inactive state prior to signaling events. The function of the CRD in autoinhibition was first indicated more than 20 years ago in studies showing that mutation of the zinc-coordinating cysteine residues within the CRAF-CRD prevented the regulatory domain from inhibiting the catalytic domain (Cutler *et al*., 1998; Winkler et al., 1998). More recently, cryoEM structures of inactive BRAF monomers further support the autoinhibitory function of the CRD, revealing that the CRD makes direct contact with both the kinase domain and a bound 14-3-3 dimer that stabilizes the autoinhibited conformation (Martinez Fiesco et al., 2021; Park et al., 2019).

Taken together, these prior studies have established the complex roles of the CRD as both a facilitator of RAS-RAF binding and RAF activation under signaling conditions and as a mediator of autoinhibition during quiescent conditions. However, the interplay between these opposing activities and their relative importance in modulating RAF function has not been well-characterized. Here, analysis of a panel of RASopathy-associated BRAF-CRD mutations provides key insights regarding BRAF regulation and reveals functional differences between RAF members that are relevant to the molecular etiology of the RASopathies.

## RESULTS

### Effect of BRAF-CRD Mutations on Phosphatidylserine Binding

RASopathy-associated mutations in the BRAF-CRD can be broadly divided into two groups – those that cluster around Loop 1 (hereon referred to as L1 mutations; Figures 1A and S1A) and those outside of Loop 1 that often introduce additional positively charged residues into the CRD (referred to as PC mutations), with Q257R being the most common CRD mutation (Figure 1A). To determine how the CRD mutations impact BRAF function, a panel of L1 (T241P, T241R, A246P) and PC (Q257R, Q262K and E275K) mutants were selected for investigation. Because a well-established property of the RAF-CRDs is their ability to interact with phosphatidylserine (PS), which contributes to the plasma membrane recruitment of RAF for kinase activation, we first examined the effect of the BRAF-CRD mutations on PS binding. For these experiments, surface plasmon resonance (SPR) was used to monitor the binding of purified WT or mutant CRD proteins (amino acids 232-284) to both nanodiscs and liposomes that contained PS (20% POPS/80% POPC). As expected, the WT-CRD demonstrated significant levels of binding to the PS-containing nanodiscs and liposomes (Figures 1B and S1B). No significant change in binding was observed for CRDs containing the L1 mutations. In contrast, CRDs containing the PC mutations displayed increased binding to the PS-containing surfaces, with the Q257R-CRD demonstrating the highest level of interaction as reflected in the normalized binding curves for the nanodisc assays and in the partition coefficients for the liposome assays (Figures 1C and 1D). As a control, binding of the CRDs to nanodiscs and liposomes that lacked PS (100% POPC) was also evaluated, and for all the CRDs, binding was reduced ∼80-97% (Figures 1B and S1B), demonstrating the preferential binding of the CRDs to lipid surfaces that contain PS. Finally, when the isolated CRD proteins were fused to GFP and their localization examined in COS-7 cells, the fusion proteins could be detected at the PS-containing cell periphery, and this localization correlated with the PS binding activity observed in the SPR analysis (Figures 1E and 1F).

### Effect of BRAF-CRD Mutations on RAF Autoinhibitory Interactions

The CRD is also known to play a role in RAF autoinhibition, which must be relieved for kinase activation to occur. To assess RAF autoinhibition, a proximity-based NanoBRET assay (Machleidt et al., 2015) was developed in which RAF autoinhibitory interactions could be quantified under live cell conditions. Previous studies have shown that when co-expressed as separate proteins, the isolated RAF regulatory (REG) domain can bind the catalytic (CAT) domain to suppress its signaling activity and that the autoinhibitory function of the REG domain requires an intact CRD (Chong and Guan, 2003; Cutler *et al*., 1998; Tran et al., 2005). Therefore, we generated expression constructs in which BRAF^REG^ (residues 1-435) contained a Halo energy acceptor tag and BRAF^CAT^ (residues 435-766) was fused to a NanoLuc energy donor tag. When co-expressed in 293FT cells, BRAF^REG^-Halo was found to interact with Nano-BRAF^CAT^, as evidenced by energy transfer and the generation of a BRET signal (Figure 2A). In addition, BRAF^REG^-Halo could suppress MEK phosphorylation induced by Nano-BRAF^CAT^ in a manner that was dose-dependent and correlated with an increased BRET signal (Figures 2A and 2B), confirming the validity of the NanoBRET assay as a measure of autoinhibitory interactions.

**Figure 2.**
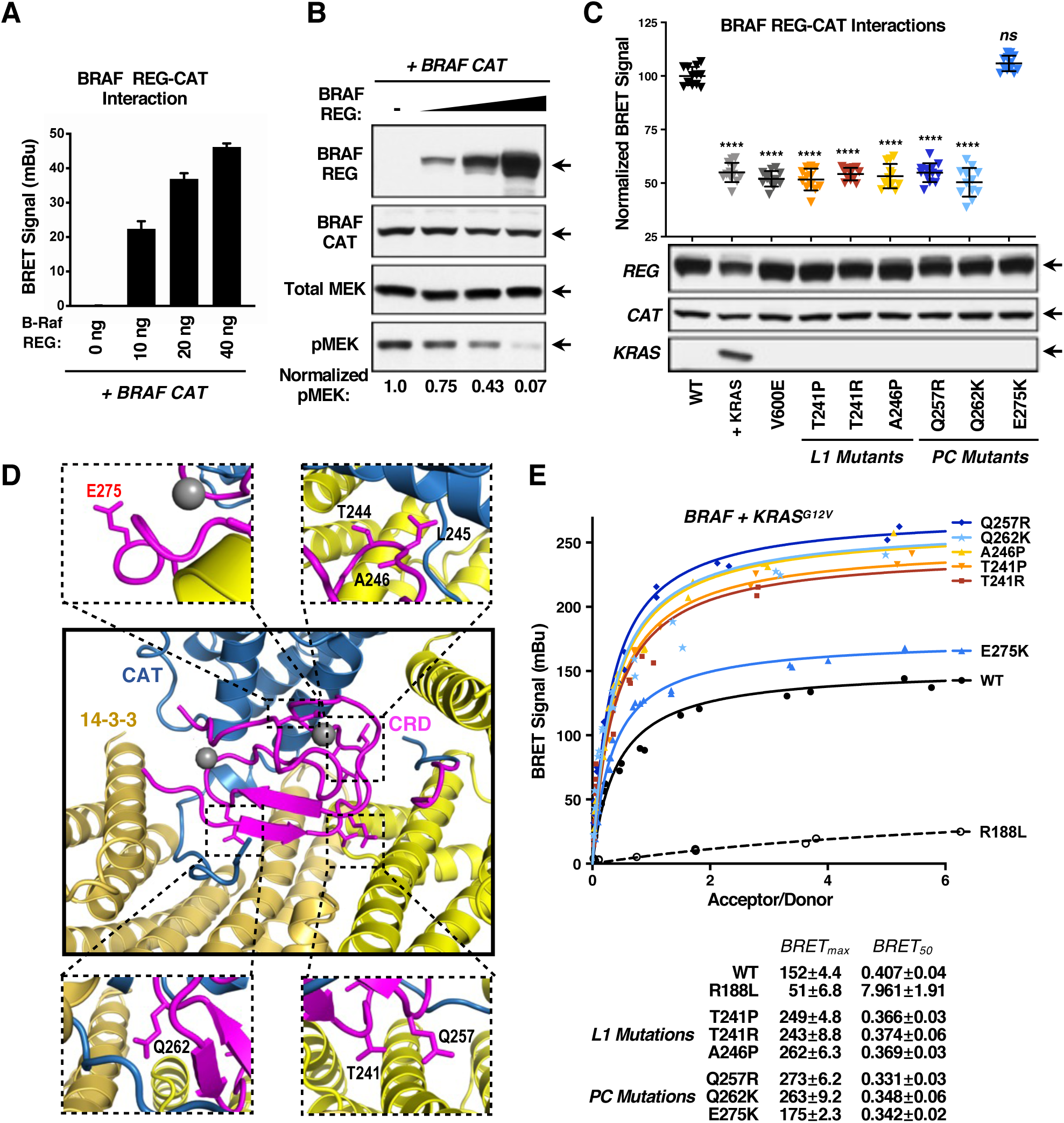
Identification of BRAF-CRD Mutations that Modulate RAF Autoinhibitory Interactions and RAS-RAF Binding in Live Cells. (A) NanoBRET interaction assay showing the BRET signal generated when increasing amounts of the WT-BRAF^REG^-Halo construct were co-transfected with a constant amount of Nano-WT-BRAF^CAT^. Bars indicate the mean of quadruplicate wells from 3 independent experiments ± SD. (B) Lysates of 293FT cells transiently expressing Nano-WT-BRAF^CAT^ alone or co-expressing Nano-WT-BRAF^CAT^ with increasing amounts of WT-BRAF^REG^-Halo were examined by immunoblot analysis for BRAF^REG^, BRAF^CAT^, and pMEK levels. Blots are representative of 3 independent experiments with similar results. (C) NanoBRET assay measuring the interaction of WT- or mutant BRAF^REG^-Halo proteins and Nano-WT-BRAF^CAT^. Also shown, is the effect of co-expressing HA-tagged KRAS^G12V^ with the WT BRET pair. Data points represent BRET signals (normalized to WT set at 100) of quadruplicate wells from 3 independent experiments ± SD. Student’s t-test (two-tailed, assuming unequal variance) was performed. ns, not significant and ****, P < 0.0001. Cell lysates were also examined by immunoblot analysis for BRAF^REG^, BRAF^CAT^, and HA-KRAS levels. (D) Cryo-EM structure of autoinhibited BRAF, showing that the CRD (magenta) contacts both the RAF catalytic domain (blue) and a bound 14-3-3 dimer (yellow). CRD residues mutated in the RASopathies are annotated (PDB: 6NYB). (E) BRET saturation curves are shown examining the interaction of WT or mutant BRAF^FL^-RLuc8 proteins with Venus-KRAS^G12V^. BRET_max_ and BRET_50_ values derived from the saturation curves are also listed ± SEM. Donor and acceptor expression levels were monitored via the RLuc8 and Venus emissions and are incorporated into the BRET calculations. Saturation curves were repeated 3 times with similar results.

The NanoBRET assay was also able to detect a relief in autoinhibitory interactions, as co-expression of a potent RAF activator, KRAS^G12V^, resulted in a 40-50% reduction in the BRET signal (Figure 2C), consistent with the binding of BRAF^REG^ to membrane-localized KRAS^G12V^ and reduced interaction with BRAF^CAT^. In addition, the NanoBRET assay was sensitive to perturbations caused by internal BRAF mutations, as incorporation of the oncogenic V600E mutation into BRAF^CAT^ reduced the BRET signal ∼50% (Figure 2C). When the various CRD mutations were introduced into BRAF^REG^, we found that all L1 mutations, along with PC mutations Q257R and Q262K, caused a significant reduction in the BRET signal, reflecting reduced binding to BRAF^CAT^ (Figure 2C). In contrast, the E275K mutation, which resulted in enhanced PS binding, had no effect on BRAF^CAT^ interactions (Figure 2C), indicating that increased PS binding alone is not sufficient to relieve autoinhibition. These findings are in agreement with the published cryo-EM structure of an autoinhibited BRAF monomer bound to dimeric 14-3-3 (Park *et al*., 2019), which shows that the PS binding interface of the CRD is occluded in the autoinhibited state and that all of the analyzed CRD mutations, with the exception of E275K, occur in or adjacent to residues that make contact with the BRAF CAT domain and/or 14-3-3 to maintain the autoinhibited conformation (Figure 2D).

### Effect of BRAF-CRD Mutations on RAS-RAF binding

Next, experiments were conducted to determine whether the CRD mutations alter the ability of BRAF to interact with activated RAS. Due to the key role of the plasma membrane in both CRD and RAS function, the RAS-RAF interaction was evaluated in the context of an intact lipid bilayer using a traditional live-cell BRET assay that had previously identified KRAS as the predominant BRAF binder (Terrell et al., 2019). For these studies, full-length BRAF tagged with Rluc8 (BRAF^FL^-RLuc8) served as the energy donor and Venus-tagged KRAS^G12V^ functioned as the energy acceptor (Venus-KRAS^G12V^). In addition, saturation curves were generated such that the BRET_max_ (the maximal level of RAS binding) and BRET_50_ (the relative affinity of the interaction) could be determined.

As expected, WT-BRAF^FL^ generated a strong BRET signal that was dramatically reduced by incorporation of the RBD R188L mutation that disrupts RAS binding (Figure 2E). Analysis of the CRD mutants revealed that proteins containing mutations that relieve autoinhibition generated a BRET_max_ signal that was increased ∼65% over that observed for WT (Figure 2E). In contrast, the E275K mutation, which enhances PS binding but does not relieve autoinhibition, resulted in a markedly weaker increase in the BRET_max_ signal (∼15%). Interestingly, when the BRET_50_ values were calculated, all of the CRD mutants, including E275K, exhibited an increase in the affinity of the KRAS interaction (lower BRET_50_ values), with the PC mutants having the lowest BRET_50_ values (Figure 2E). These findings suggest that the PS binding activity of the BRAF-CRD may impact the affinity of KRAS binding. Indeed, when the BRET assays were conducted using the isolated BRAF^REG^ domain, where autoinhibition would not be a factor, the CRD L1 mutants generated BRET signals that were equivalent to WT-BRAF^REG^ in terms of both the BRET_max_ and BRET_50_ values (Figure S2), consistent with these mutations only altering autoinhibition. In contrast, an elevated level and affinity of RAS binding was observed for the PC mutants that correlated with the effect of these mutations on PS binding (Q257R>E275K>Q262K; Figure S2).

### All BRAF-CRD Mutants Require RAF Dimerization for Their Enhanced Signaling Activity but Differ in Their Dependence on RAS Binding

The above results indicate that the BRAF-CRD mutations vary in their effects on RAF autoinhibition as well as PS and RAS binding, with the L1 mutations causing a relief in autoinhibition (which we refer to as Class A; Table S1), the PC Q257R and Q262K mutations both relieving autoinhibition and increasing PS binding (Class B), and the PC E275K mutation only causing enhanced PS binding (Class C). Moreover, relief of autoinhibition alone was sufficient to increase the overall levels of RAS-RAF binding in live cells, whereas enhanced PS binding correlated with an increase in the affinity of the interaction, either by stabilizing membrane interactions and/or facilitating direct CRD-RAS contact. To determine how these differing effects influence the signaling activity of BRAF, full-length BRAF proteins containing the CRD mutations were evaluated in focus forming assays using NIH-3T3 cells. In contrast to WT-BRAF^FL^, all of the CRD mutants were able to induce focus formation (Figures 3A and S3A), with proteins containing mutations that relieve autoinhibition (Class A and B) being significantly more transforming than the E275K mutant (Class C), which has unimpaired autoinhibitory activity. The Class A and B mutants were also more effective at promoting MEK phosphorylation in NIH-3T3 cells than was E275K-BRAF^FL^ (Figure 3B).

**Figure 3.**
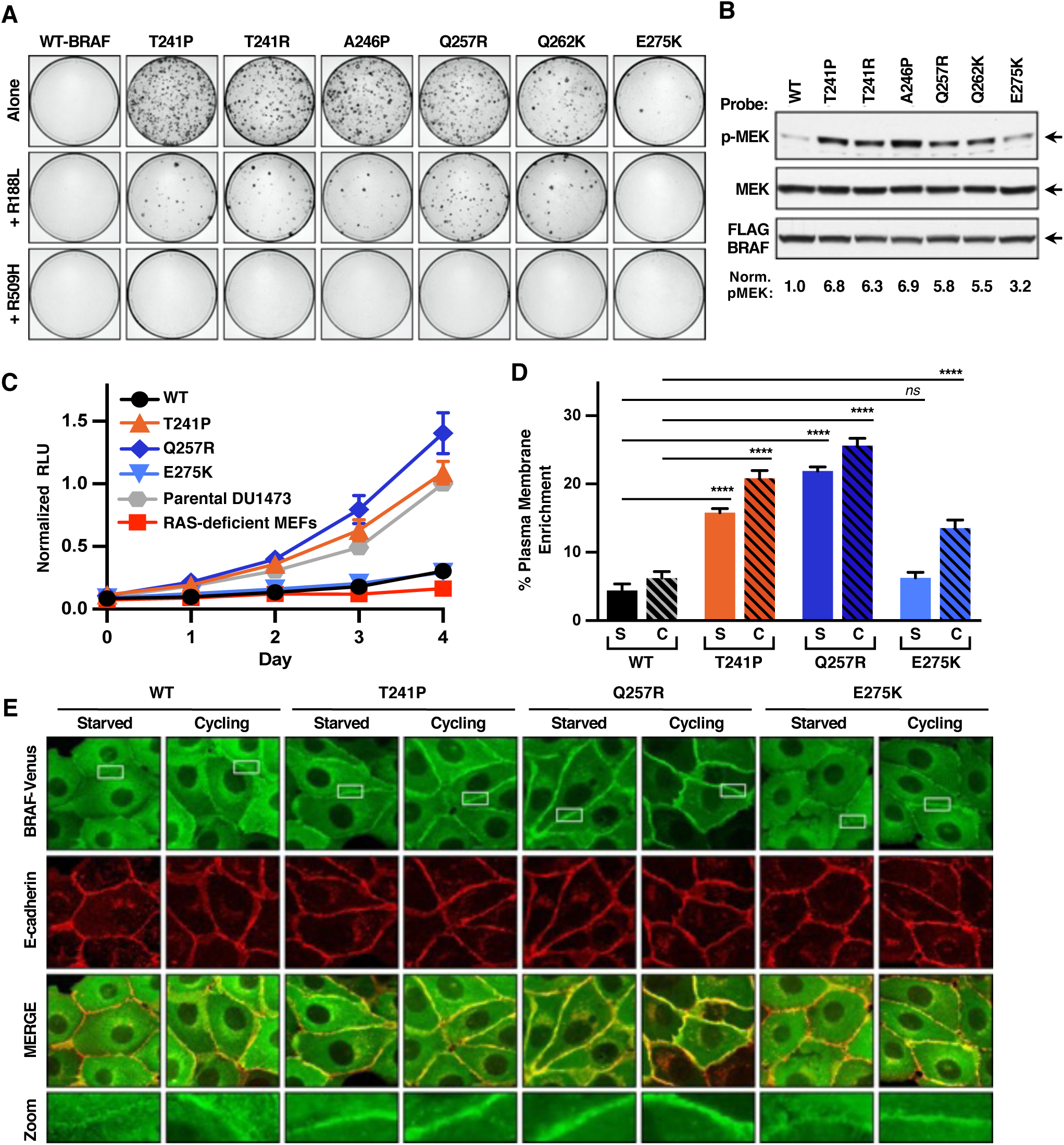
Effect of BRAF-CRD Mutations on the Signaling Activity and Plasma Membrane Localization of BRAF. (A) NIH-3T3 cells were infected with retroviruses encoding the indicated WT or mutant FLAG-BRAF^FL^ proteins. Three weeks post-infection, foci were visualized by methylene blue staining. Assays were repeated 3 times with similar results. (B) Lysates of serum-starved NIH-3T3 cells expressing the indicated FLAG-BRAF^FL^ proteins were examined for pMEK, FLAG-BRAF^FL^, and total MEK levels. Blots are representative of 3 independent experiments with similar results. (C) The proliferation of RAS-deficient MEF lines expressing the indicated FLAG-BRAF^FL^ proteins was monitored for 4 days using Cell TiterGlo. Data points represent the mean of quadruplicate wells from 2 independent experiments ± SEM. (D and E) MDCK cells stably expressing the indicated BRAF^FL^-Venus variants were left in growth media or serum-starved for 18 hrs, following which they were fixed, stained for E-cadherin, and then imaged. Percent plasma membrane enrichment of BRAF^FL^ (D) was calculated from fluorescence traces of n=20 cells ± SEM that were imaged by confocal microscopy (E). Imaging studies were repeated 3 times with similar results.

Additional studies revealed that the focus forming activity of the CRD mutants was dependent on RAF dimerization as incorporation of the R509H mutation to disrupt dimerization prevented foci formation in all cases (Figures 3A and S3A). However, the importance of RAS binding appeared more variable in that when the RBD R188L mutation was introduced, proteins containing the Class A and B mutations, which disrupt autoinhibition, retained 20-40% of their focus forming activity even in the absence of an intact RBD, with Q257R/R188L-BRAF^FL^ exhibiting the highest level of activity. In contrast, when the R188L mutation was introduced into the BRAF protein containing the Class C E275K mutation, which only enhances PS binding, no foci were observed (Figure 3A).

To further assess the contribution of RAS to the signaling activity of the CRD variants, cell proliferation studies were conducted using RAS-deficient mouse embryo fibroblasts (MEFs) that stably expressed WT-BRAF or a representative of each of the CRD mutant classes: T241P (Class A), Q257R (Class B), or E275K (Class C) (Figure S3B). As shown in Figure 3C, RAS-deficient MEFs overexpressing WT-BRAF^FL^ or the E275K mutant displayed minimal proliferation. In contrast, cells expressing T241P- or Q257R-BRAF^FL^, whose mutations relieve autoinhibition, exhibited proliferation rates similar to or higher than that observed for the parental KRAS-expressing DU1473 MEF line (Figure 3C), suggesting that relief of autoinhibition allows these mutants to promote cell growth in the absence of canonical RAS proteins.

Next, we examined how the different CRD mutation classes affect the subcellular localization of BRAF. For these studies, stable MDCK lines that express WT, T241P (Class A), Q257R (Class B), or E275K (Class C) BRAF^FL^-Venus-tagged proteins (Figure S3C) were generated and examined under serum-starved or cycling growth conditions. The cells were imaged via confocal microscopy, and the plasma membrane enrichment of the various BRAF^FL^-Venus proteins was determined using E-cadherin as a plasma membrane marker. Under serum-starved conditions, where RAS-GTP levels would be low, WT-BRAF^FL^ exhibited primarily a perinuclear/cytosolic localization with minimal levels detected at the plasma membrane (Figures 3D and 3E). A similar localization was observed for E275K-BRAF^FL^, whose mutation only enhances PS binding (Figure 3D and 3E). However, T241P- and Q257R-BRAF^FL^, whose mutations relieve autoinhibition, displayed a 3.6 and a 5-fold enrichment at the plasma membrane, respectively, when compared to WT-BRAF^FL^. Interestingly, under cycling conditions where RAS-GTP levels would be elevated, the plasma membrane localization of all the CRD mutants, including E275K-BRAF^FL^, was significantly increased over WT levels (Figures 3D and 3E). Taken together, the above data suggest that relief of CRD-mediated autoinhibition alone is sufficient to increase the pool of BRAF at the plasma membrane to levels needed for cell signaling and is similar to the RAS-independent behavior observed for membrane-localized Raf-CAAX proteins (Leevers et al., 1994; Stokoe et al., 1994). Moreover, these findings demonstrate the RAS-dependency of the E275K mutant in requiring the interaction with RAS to disrupt the autoinhibited state, and then the role of its enhanced PS binding activity, once autoinhibition is released.

### BRAF-CRD Mutations Increase the Biological Activity of BRAF in Zebrafish Embryos

To investigate the effects of the CRD mutations on BRAF function in a developing organism, the zebrafish model system was used, as expression of RASopathy mutants has been shown to induce defects in zebrafish embryos similar to those observed in RASopathy patients, including alterations in the heart and craniofacial structures (Patterson and Burdine, 2020). In addition, monitoring the effects of RASopathy mutants on early convergent-extension cell movements, which results in elongated embryos with an abnormal axis ratio, has been used to assess mutation severity (Jindal et al., 2017). As with the above experiments, WT-BRAF and a representative of each of the three CRD mutant classes was examined (Class A T241P, Class B Q257R, and Class C E257K). In convergent extension assays, expression of T241P- or Q257R-BRAF had a more significant effect on the embryo axis ratio than did E275K-BRAF, resulting in an axis ratio of 1.14 and 1.13 for the Class A and B mutants, respectively, versus 1.05 for the E275K Class C mutant (Figure 4A). In addition, when embryos were examined 3 days post-fertilization 26-30% of embryos expressing the Q257R or T241P mutants developed heart edema, whereas less than 10% of those expressing E275K-BRAF displayed this phenotype (Figure 4B). Finally, the effect of the CRD mutants on craniofacial structures was determined by measuring the angle of the ceratohyal, which corresponds to the width and bluntness of the head and is an excellent marker for hypertelorism, a common feature of RASopathy patients. Using the transgenic *Tg (col2a1aBAC:GFP)* (Askary et al., 2015) zebrafish line to visualize cartilage cells, we found that expression of either the T241P or Q257R mutant caused a significant increase in the angle of the ceratohyal, whereas expression of the E257K mutant did not (Figure 4C). These embryo assays and the above signaling experiments indicate that E275K is a weaker gain-of-function mutation, a finding consistent with the phenotypic analysis of a CFC patient possessing this mutation and who does not exhibit characteristics frequently observed in CFC patients with BRAF-CRD mutations, such as hypertelorism, short stature, or mental retardation (Sarkozy et al., 2009) (Figure 4D and Tables S2). Notably, these findings further suggest that relief of CRD-mediated autoinhibition is the primary factor determining the severity of the BRAF-CRD mutations.

**Figure 4.**
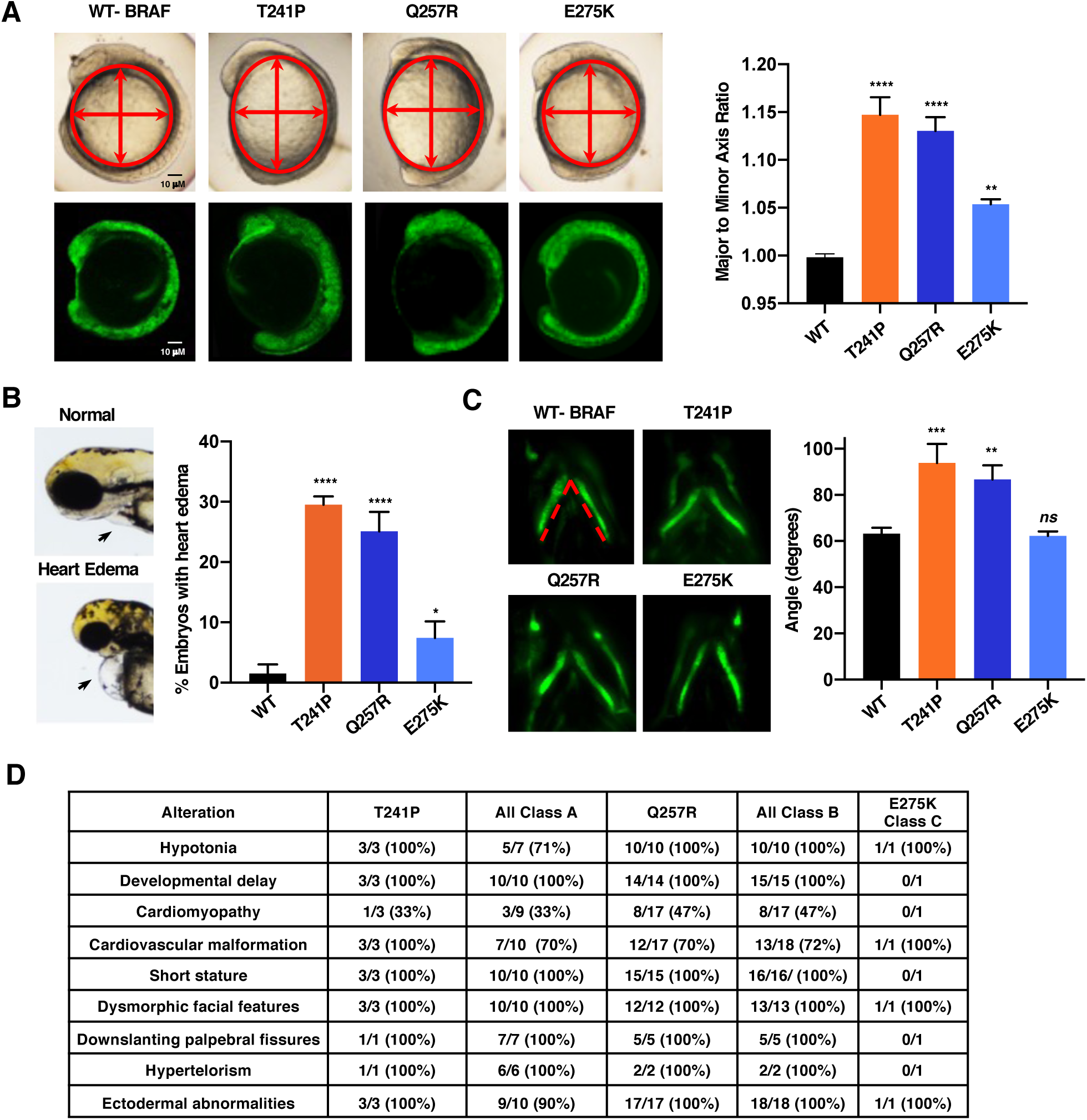
Expression of BRAF-CRD Mutants Induces Developmental Alterations in Zebrafish Embryos. (A) mRNA encoding the indicated Venus-tagged BRAF proteins were injected into one-cell stage zebrafish embryos and the embryos were measured at 11 hrs post fertilization (hpf) to determine the major to minor axis ratio. Shown are representative images of the embryos and the expression levels of the BRAF-Venus proteins (left). The average axis ratio of embryos analyzed in 3 independent experiments is shown ± SEM (right). Average axis ratio of uninjected embryos is 1 ± SEM 0.005. (B) Embryos injected as in (A) and were examined 3 days post fertilization (dpf) for heart edema. Zebrafish embryos with normal and edemic hearts are shown (left). The graph represents the percentage of embryos with heart edema in 3 independent experiments ± SEM (right). (C) One-cell stage embryos from the *Tg (col2a1aBAC:GFP)* transgenic line were injected as in (A). At 5dpf, the ceratohyal was marked by GFP, and the ceratohyal angle was determined (dotted red line). Representative images are shown (left). The graph represents the quantification of the angle of the ceratohyal from 4 independent experiments ± SEM. Average ceratohyal angle for uninjected embryos is 59° ± SEM 3.0. (D) Summary of phenotypic characteristics of RASopathy patients with BRAF-CRD mutations. Data were collected from <u>nseuronet.com</u> and references are cited in Table S2. For statistical analysis (A-C), one-way analysis of variance (ANOVA) with Bonferroni correction was used. ns, not significant; *, *P* < 0.05, P **, *P* < 0.01; ***, *P* < 0.001, and ****, *P* < 0.0001.

### The CRDs of BRAF and CRAF Differ in Their Ability to Bind PS

The CRD is a hotspot for RASopathy-associated mutations in BRAF but not CRAF, despite reports that the CRAF-CRD also plays a role in autoinhibition as well as PS and RAS binding (Cutler *et al*., 1998; Fang *et al*., 2020; Improta-Brears et al., 1999; Tran *et al*., 2021; Travers et al., 2018; Winkler *et al*., 1998). Therefore, studies were initiated to compare the relative ability of the BRAF- and CRAF-CRDs to perform these functions. Through SPR analysis, we found that the CRAF-CRD exhibited specific binding to PS-containing nanodiscs; however, the level of binding was ∼30% of that observed for the BRAF-CRD (Figures 5A, 5B and S4). Consistent with the SPR analysis, when the isolated CRD proteins were fused to GFP and their localization examined in COS-7 cells, the BRAF-CRD was clearly visible at the cell periphery, whereas the isolated CRAF-CRD was observed primarily in the cytosol and nucleus (Figure 5C). Of note, the nuclear localization of the GFP-CRD is likely due to its small size (33.8 kDa), which would permit diffusion into the nucleus as has been reported for the PKC CRD (Oancea et al., 1998; Verbeek et al., 2008).

**Figure 5.**
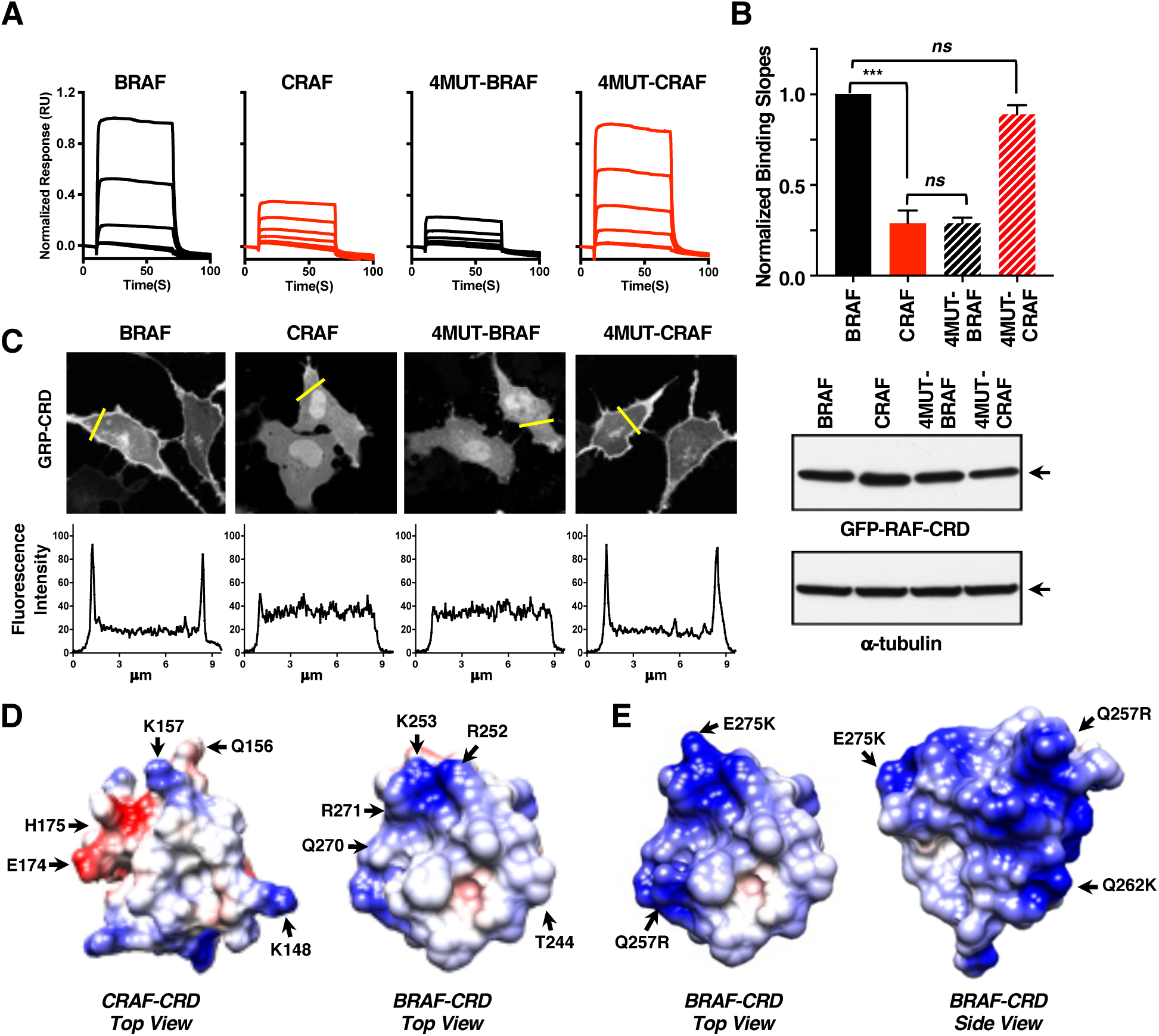
The CRDs of BRAF and CRAF Differ in Their Abilities to Bind PS. (A) The indicated BRAF- and CRAF-CRD proteins were examined by SPR for binding to PS-containing nanodiscs. Sensorgrams were normalized to the WT-BRAF-CRD binding response. (B) Binding slopes for the CRAF-CRD and the indicated CRD mutants were normalized to the binding slope of the BRAF-CRD (with BRAF-CRD equaling 1), and the graph represents the mean of 3 independent experiments ± SD. Student’s *t*-test (two-tailed, assuming unequal variance) was performed. ns, not significant and ***, *P* < 0.001. (C) Serum-starved COS-7 cells transiently expressing the indicated GFP-CRD proteins were examined by immunoblot analysis and confocal microscopy. A tracing depicting the GFP intensity in the area indicated by the yellow line is shown below the images. The images are representative of 2 independent experiments with similar results. (D) Electrostatic surface representation of the BRAF-CRD (PDB: 6NYB) and CRAF-CRD (PDB: 1FAR) structures are shown, with blue and red representing positively and negatively charged areas, respectively, and key residues indicated. (E) Electrostatic surface charge distribution of the BRAF-CRD (PDB: 6NYB) is shown, with the PC mutations indicated.

To ascertain the residues responsible for the differing PS binding activities of the BRAF and CRAF-CRDs, we examined the subcellular localization of a series of GFP-tagged CRD proteins in which non-conserved residues between the BRAF and CRAF-CRDs had been exchanged. From this analysis, four BRAF residues (F250, R252, Q270, and R271) were identified that when substituted for the equivalent residues in the CRAF-CRD (4MUT-CRAF, I154F Q156R, E174Q, and H175R) resulted in an enhanced cell surface localization that was comparable to the BRAF-CRD (Figure 5C). Likewise, when the equivalent residues of CRAF were swapped into the BRAF-CRD (4MUT-BRAF), the 4MUT-BRAF-CRD localized in a manner similar to the CRAF-CRD (Figure 5C). SPR analysis confirmed that these localizations correlated with the PS binding activity of the CRDs and that the PS binding activity of the BRAF- and CRAF-CRD could largely be reversed by exchanging these four non-conserved residues (Figures 5A-C and S4B). Interestingly, surface charge comparisons of the RAF-CRD structures revealed that the four non-conserved residues determining the affinity of PS binding localize to the top surface of the CRD that contains the hydrophobic pocket formed by the L1 loop (Figure 5D). The CRAF residue E174 adds an acidic amino acid to this region, which would be repulsive in the context of the negatively charged plasma membrane, whereas the BRAF residues R252 and R271, together with the conserved lysine at position 253 (K253), form a positively charged patch that would be conducive for increased binding to the negatively charged head groups of PS. Moreover, the RASopathy PC mutations would place additional basic residues adjacent to (Q257R and E275K) or slightly below (Q262K) this surface (Figure 5E).

### The BRAF-CRD is a More Potent Mediator of Autoinhibition Than is the CRAF-CRD

To compare the autoinhibitory activities of the BRAF and CRAF-CRDs, a BRAF^REG^ construct in which the BRAF-CRD was replaced with that of CRAF was generated, following which the ability of the C-CRD-BRAF^REG^ to interact with the BRAF^CAT^ domain was monitored using the NanoBRET assay. In comparison to BRAF^REG^, the BRET signal generated by C-CRD-BRAF^REG^ was significantly lower, indicating a reduced ability to mediate autoinhibitory interactions (Figure 6A). When similar constructs were generated to evaluate CRAF autoinhibition, an interaction between the CRAF^REG^ and CRAF^CAT^ domains was observed, but the BRET signal generated was lower than that observed for the BRAF domains (Figure 6A). However, when the CRAF^REG^ construct contained the BRAF-CRD, the BRET signal increased approximately 40%. In addition, when the BRAF^REG^ proteins were evaluated for binding to CRAF^CAT^ and vice versa, similar results were observed (Figure 6A). These findings confirm that the BRAF-CRD is a more potent mediator of autoinhibition and that this increased activity is not determined by differences in the BRAF versus CRAF catalytic domains.

**Figure 6.**
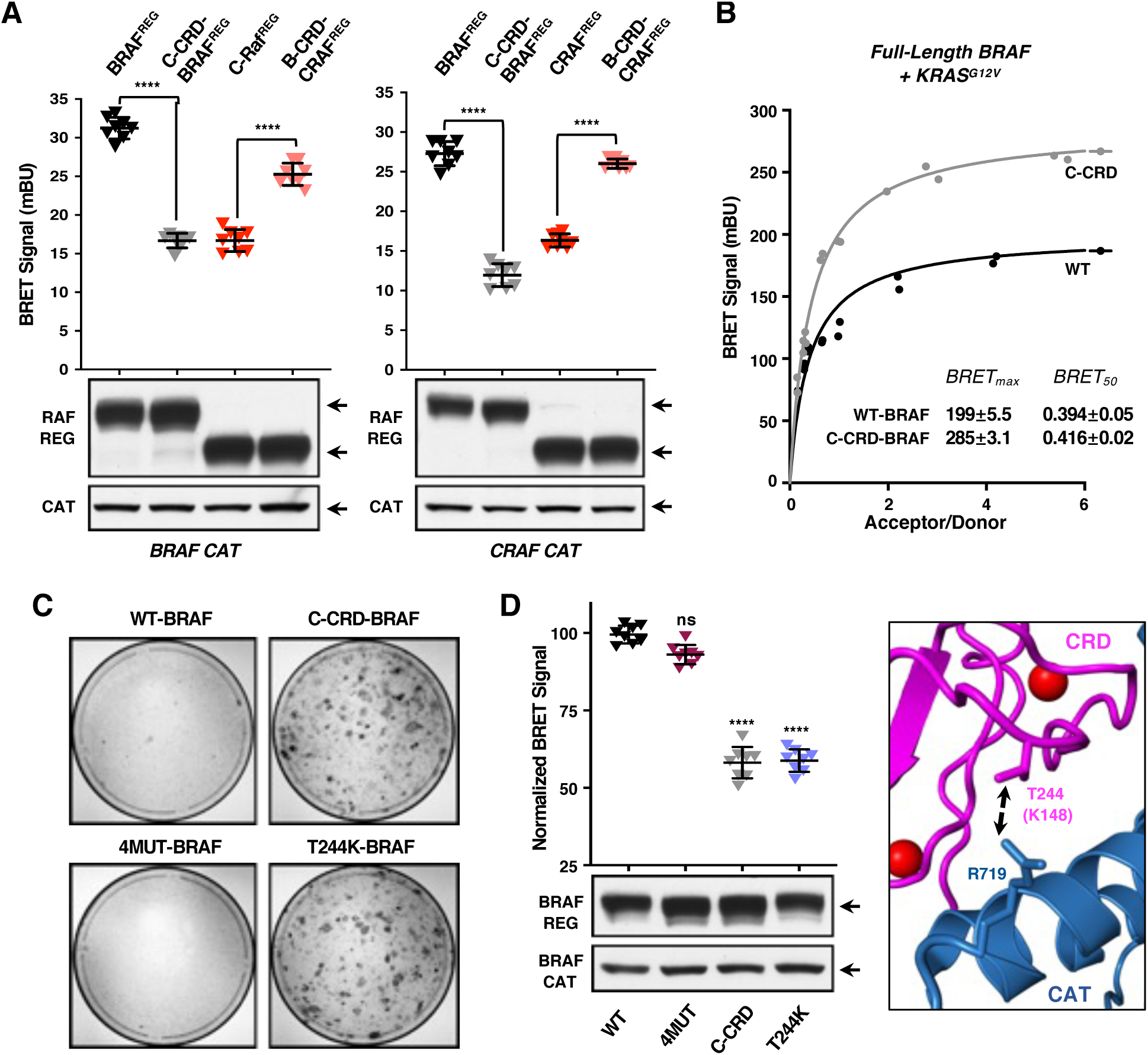
The BRAF- and CRAF-CRDs Differ in Their Abilities to Mediate Autoinhibition. (A) NanoBRET interaction assay showing the BRET signal generated when the indicated RAF^REG^-Halo constructs were co-transfected with Nano-WT-BRAF^CAT^ (left) or Nano-WT-CRAF^CAT^ (right). Cell lysates were also examined by immunoblot analysis for RAF^REG^ and RAF^CAT^ proteins levels. Data points represent quadruplicate wells from 3 independent experiments ± SD. (B) BRET saturation curves are shown examining the interaction of WT or C-CRD-BRAF^FL^-RLuc8 proteins with Venus-KRAS^G12V^. BRET_max_ and BRET_50_ values are listed ± SEM. Saturation curves were repeated 3 times with similar results. (C) NIH-3T3 cells were infected with retroviruses encoding the indicated FLAG-BRAF^FL^ variants. Three weeks post-infection, foci were visualized by methylene blue staining. Assays were repeated 2 times with similar results. (D) NanoBRET assay measuring the interaction of WT- or mutant BRAF^REG^-Halo proteins and Nano-WT-BRAF^CAT^. Cell lysates were also examined by immunoblot analysis for BRAF^REG^ and BRAF^CAT^ proteins levels. Data points represent quadruplicate wells from 3 independent experiments ± SD. Proximity of the BRAF-CRD residue T244 (magenta) and the BRAF catalytic domain residue R719 (blue) is also shown (PDB: 6NYB). For (A, D), student’s *t*-test (two-tailed, assuming unequal variance) was performed. ns, not significant and ****, *P* < 0.0001.

Next, we examined the effect of exchanging the CRAF-CRD for that of BRAF on binding between full-length BRAF and activated KRAS using our traditional BRET assay. Consistent with the lower PS binding activity of the CRAF-CRD, the affinity of the RAS-RAF interaction was reduced for C-CRD-BRAF^FL^, whereas the overall levels of RAS-RAF binding were higher than that observed for WT-BRAF^FL^ and correlated with the reduced autoinhibitory activity of the CRAF-CRD (Figure 6B). Strikingly, in focus forming assays, C-CRD-BRAF^FL^, but not WT-BRAF^FL^, was able to transform NIH-3T3 cells (Figure 6C), thus demonstrating the critical role of the BRAF-CRD in maintaining BRAF in an autoinhibited, non-signaling state.

Because both the PS binding and autoinhibitory activities of the BRAF-CRD were elevated in comparison to the CRAF-CRD, we next examined whether the four non-conserved BRAF residues identified to mediate increased PS binding might also be responsible for the elevated autoinhibitory activity. Using the NanoBRET assay, 4MUT-BRAF^REG^ showed no impairment in mediating autoinhibitory interactions with BRAF^CAT^ (Figure 6D), indicating that these residues are not critical for autoinhibition. Through the analysis of a series of BRAF^REG^ proteins in which non-conserved residues between the BRAF and CRAF-CRDs were exchanged, a single residue swap in L1, T244K (K149 in CRAF), was found to reduce interactions between the BRAF^REG^ and BRAF^CAT^ domains to a level observed with C-CRD-BRAF^REG^ (Figure 6D; Table S3). Interestingly, in the autoinhibited structure of the BRAF monomer, T244 lies in close proximity to R719 in the BRAF catalytic domain (Martinez Fiesco *et al*., 2021; Park *et al*., 2019) and placement of a lysine residue at position 244 would likely generate repulsive forces with R719, thereby destabilizing the autoinhibitory contacts of the CRD (Figure 6D). Finally, when the effect of the T244K substitution was examined in focus forming assays, we found that T244K-BRAF^FL^, but not 4MUT-BRAF^FL^, was as transforming as C-CRD BRAF^FL^ (Figure 6C).

### Analysis of CRAF Proteins Containing Mutations Analogous to the RASopathy BRAF-CRD Mutations

The basal catalytic activity of BRAF is significantly higher than that of CRAF, due primarily to the constitutive negative charge of a region in the BRAF kinase domain known as the negative charge regulatory region or N-region (residues 444-449; (Emuss et al., 2005; Marais et al., 1997). Therefore, it is possible that the enhanced autoinhibitory activity of the BRAF-CRD may be specifically required to quell the high basal catalytic activity of BRAF and may explain why the CRAF-CRD is not a mutational hotspot in the RASopathies. To investigate this possibility, constructs expressing full-length CRAF that contained CRD mutations equivalent to T241P, Q257R, and Q262K (CRAF T145P and N161R, and Q166K) were generated and examined for their focus forming potential. As shown in Figure 7A, none of these mutants could induce foci. However, when the N-region of these CRAF mutants was changed to that of BRAF, conferring high basal catalytic activity (Figure 7B), the CRAF-CRD mutants were now transforming (Figure 7A). Moreover, CRAF proteins containing only the BRAF N-region sequences were also weakly transforming, consistent with elevated catalytic activity of the hybrid protein and the reduced autoinhibitory activity of the CRAF-CRD. These findings suggest that the CRD and autoinhibition play a more important role in BRAF regulation, functioning to maintain BRAF in an inactive state during non-signaling conditions.

**Figure 7.**
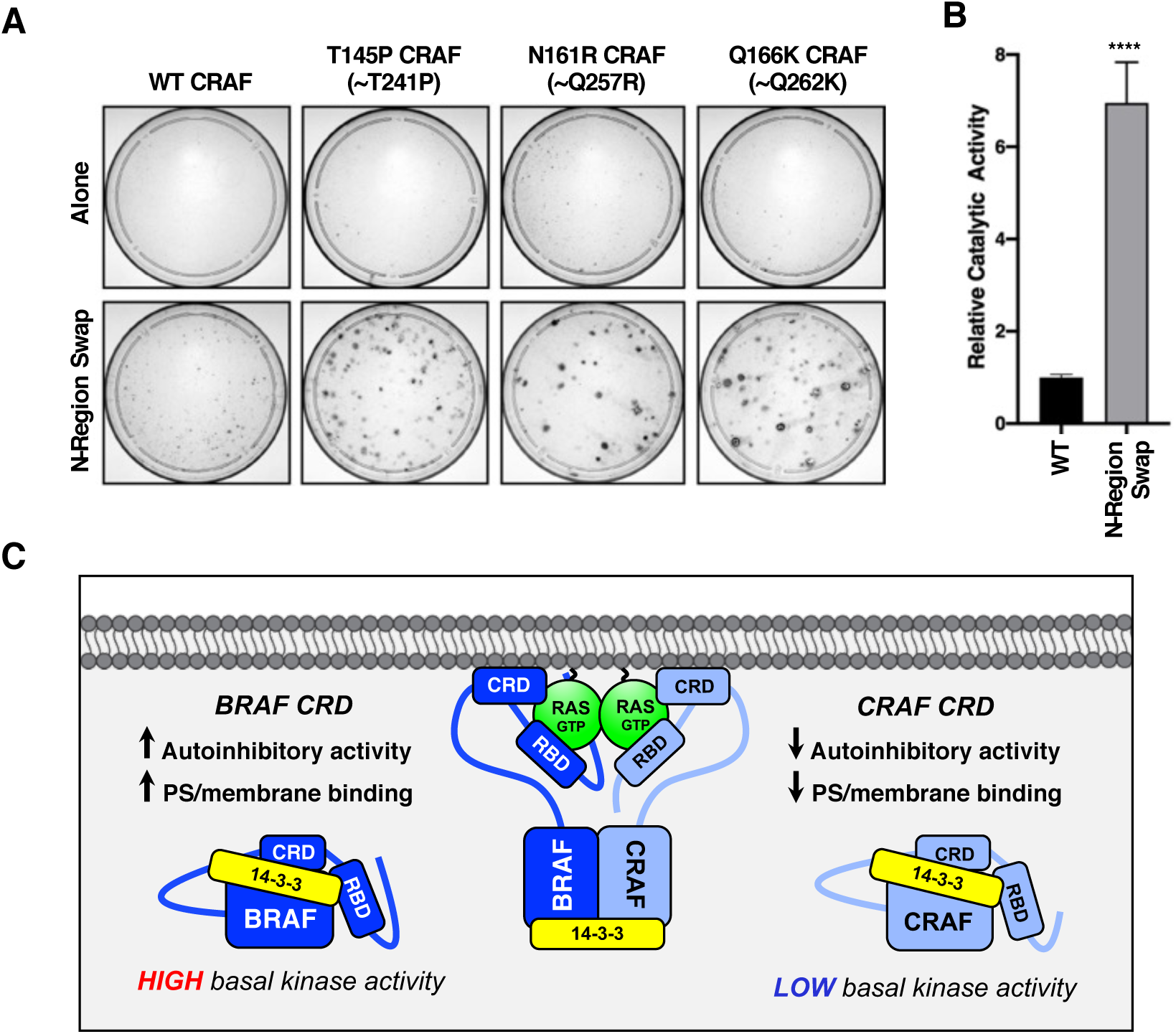
Analysis of CRAF Proteins Containing Mutations Analogous to RASopathy-associated BRAF-CRD Mutations. (A) NIH-3T3 cells were infected with retroviruses encoding the indicated WT or mutant FLAG-CRAF^FL^ proteins. Three weeks post-infection, foci were visualized by methylene blue staining. Assays were repeated three times with similar results. (B) In vitro kinase assays were performed to demonstrate the increased catalytic activity of FLAG-CRAF^FL^ containing the N-region sequence of BRAF (N-region swap) versus WT-FLAG-CRAF^FL^ (with WT activity equaling 1). Data represent the mean of duplicate samples from 3 independent experiments ± SD. Student’s *t*-test (two-tailed, assuming unequal variance) was performed. ****, *P* < 0.0001. (C) Model comparing the properties of the BRAF- and CRAF-CRDs. See text for details.

## DISCUSSION

Early investigation of the RAF kinases demonstrated the role of the CRD in PS binding, the RAS-RAF interaction, and autoinhibition; however, the relative interplay between these activities and their importance for RAF regulation and function have not been well explored. In this study, we analyzed a panel of RASopathy-associated BRAF-CRD mutations for their effect on these known activities of the CRD using a variety of approaches, including the use of a BRET-based assay to quantify RAF autoinhibitory interactions under live cell conditions. In addition, we examined the impact of these mutations on the biological function of BRAF in cell culture systems and zebrafish embryos, and we compared the properties of the BRAF-CRD with those of the CRAF-CRD. Through this analysis, we find that the BRAF-CRD is a stronger mediator of both PS binding and autoinhibition than is the CRAF-CRD and that the RASopathy-associated BRAF-CRD mutations can be classified into three functional groups: Class A, causing relief in autoinhibition; Class B, relieving autoinhibition and enhancing PS/plasma membrane binding; and Class C, enhancing PS/membrane binding alone.

Strikingly, the ability of the mutations to alter distinct activities of the CRD was found to have direct consequences on BRAF subcellular localization, RAS binding properties, and the degree of RAS-dependency for signaling activity. For example, analysis of the BRAF-CRD mutants revealed that relief of autoinhibition alone was sufficient to robustly increase the overall levels of RAS-RAF binding, whereas enhanced CRD-PS binding contributed to the affinity of the RAS-RAF interaction, either through membrane interactions or facilitating direct RAS contact. In addition, the CRD mutants with impaired autoinhibitory activity (Class A and B) displayed an increased presence at the plasma membrane even in serum-starved conditions and exhibited a level of RAS-independent activity, retaining 20-40% of their focus forming ability in the absence of an intact RBD and demonstrating the ability to drive the proliferation of RAS-deficient MEFs. In contrast, the Class C E257K mutant, which is unimpaired for autoinhibition, was fully dependent on RAS for its signaling activity in that no foci were induced when the R188L mutation was added and that this mutant, like WT-BRAF, promoted minimal growth of RAS-deficient MEFs. Moreover, in serum-starved cells, both the WT and E275K-BRAF proteins were found primarily in the cytoplasm, and only under cycling conditions where RAS-GTP levels would be elevated, did E275K-BRAF exhibit a plasma membrane localization that was significantly increased over WT, consistent with the enhanced PS binding activity of this mutant.

With regard to the analysis of the CRD mutant proteins in the zebrafish model system, the Class A and B mutants were also found to cause more pronounced developmental defects than did the Class C E275K mutant when expressed in embryos. It is interesting to note that the evaluation of one RASopathy patient with the Class C E257K mutation has been reported, and this patient does not display several attributes reported for all patients possessing Class A T241P and Class B Q257R mutations, such as hypertelorism, short stature, and development delays (Sarkozy *et al*., 2009) (Table S2), suggesting that relief of autoinhibition is the primary factor determining the phenotypic severity of these mutations. Although the number of patient reports available is quite limited, the NanoBRET assay to measure RAF autoinhibition and the functional assays described here may be particularly useful in predicting the severity of new BRAF-CRD mutations as they are identified through various ongoing and future genomic sequencing projects.

In terms of BRAF regulation, our findings indicate that for WT-BRAF and the Class C E275K mutant, RAS binding is needed to disrupt the autoinhibited state, which in turn, would expose the ligand binding interface of the CRD for contact with the plasma membrane and RAS, a model supported by a recent cryo-EM structure showing that in autoinhibited BRAF:14-3-3_2_ complexes, the RAS binding interface of the RBD is largely exposed, whereas the ligand binding interface of the CRD is occluded (Martinez Fiesco *et al*., 2021). In contrast, the Class A and B mutations relieve CRD-mediated autoinhibition themselves, and this property together with the high PS binding activity of the BRAF-CRD, allows these mutants to constitutively localize to the plasma membrane at a level that may facilitate self-dimerization and kinase activation, similar to the RAS-independent behavior observed for membrane-localized RAF-CAAX proteins (Leevers et al., 1994; Stokoe et al., 1994). This model also provides an explanation for why the Q257R mutant, whose mutation relieves autoinhibition and causes the largest increase in PS and plasma membrane binding, exhibited the highest level of RAS-independent activity. However, despite its enhanced RAS-independent activity, when RAS was present, the signaling and biological activity of Q257R-BRAF was not elevated in comparison to the Class A mutants, which only have impaired autoinhibitory activity. Insight as to why this might be the case comes from published structures of the CRAF RBD-CRD bound to KRAS and modeling of this complex at the plasma membrane (Tran *et al*., 2021; Travers et al., 2020; Travers *et al*., 2018). More specifically, these studies suggest that the CRD may play a more critical role in RAS-dependent RAF activation by helping orient the RAS-RAF complex at the plasma membrane for productive dimer formation rather than in facilitating complex formation itself. Further supporting this idea, specific mutations in the PS/membrane binding region or the RAS binding interface of the CRAF-CRD have been found to have a significant effect on RAF kinase output, yet have little impact on RAS-RAF binding (Daub *et al*., 1998; Tran *et al*., 2021). In the case of Q257R-BRAF, this mutant would be predicted to form additional interactions with the PS head groups, potentially altering the position of the CRD with respect to the membrane and RAS as well as the overall orientation of the RAS-RAF complex, which could have a detrimental effect on RAF activity.

In this study, we also compared the properties of the BRAF- and CRAF-CRDs (Figure 7C) and found that the BRAF-CRD was a stronger mediator of PS-binding and autoinhibition and that exchanging the BRAF-CRD with that of CRAF was sufficient to activate the transforming potential of BRAF. Our findings further indicate further indicate that the enhanced autoinhibitory activity of the BRAF-CRD is required to maintain BRAF in a quiescent state, as BRAF has the highest basal catalytic activity of all the RAF kinases, due largely to the constitutive negative charge of the BRAF N-region sequences. In support of this model, CRAF proteins containing the BRAF N-region exhibited significantly increased catalytic activity and were found to be weakly transforming, consistent with the reduced autoinhibitory activity of the CRAF-CRD. Moreover, while incorporation of mutations analogous to the RASopathy-associated T241P, Q257R, or Q262K into the CRAF-CRD was not sufficient to promote cellular transformation, their incorporation into CRAF proteins containing the BRAF N-region resulted in mutant proteins that had significant transforming activity. Thus, the high basal catalytic activity of BRAF combined with the critical autoinhibitory function of the BRAF-CRD may account for why the CRD is a mutational hotspot for BRAF but not CRAF in the RASopathies. However, once autoinhibition is relieved, the enhanced PS binding activity of BRAF-CRD may aid in keeping BRAF at the plasma membrane, given the lower RAS-binding affinity of BRAF versus CRAF in live cells (Terrell *et al*., 2019).

In conclusion, analysis of the RASopathy BRAF-CRD mutations has provided significant insight regarding the relative importance of the distinct activities of the BRAF-CRD and has revealed functional differences between RAF family members that are relevant to human disease (Fig. 7C). Notably, this study also demonstrates the essential role of the CRD in maintaining BRAF autoinhibition, which if relieved, results in constitutive plasma membrane localization and increased RAS-dependent and RAS-independent function, further suggesting that stabilizing the autoinhibited state using pharmacological approaches may have therapeutic benefit in preventing the aberrant activation of BRAF that frequently occurs in human cancer and the RASopathies.

## ACKNOWLEDGMENTS

We thank John-Paul Denson, Matt Drew, Dominic Esposito, Bill Gillette, Jose Sanchez Hernandez, Shelley Perkins, Nitya Ramakrishnan, Kelly Snead, Troy Taylor, Vanessa Wall, and Timothy Waybright (FNLCR Protein Expression Laboratory) for the production and characterization of the isolated CRD proteins for SPR analysis, and Linda Miller (LCDS) for technical support. This project has been funded in whole or in part with Federal funds from the National Cancer Institute, National Institutes of Health, under project number ZIA BC 010329 (D.K.M) and Contract No. HHSN261200800001E (A.G.S.).

## AUTHOR CONTRIBUTIONS

R.S.-S. and D.K.M. conceived the project, designed the experiments, and interpreted the data with contributions from A.G.S. R.S.-S. developed the NanoBRET assay to quantify RAF autoinhibitory interactions and conducted all assays, E.M.T. generated critical reagents and performed the RAS-RAF BRET interaction assays; E.M.T., D.A.R., and R.S.-S. conducted the transformation assays; C.A. and M.E.W. performed the surface plasmon resonance analysis; C.I and J.S performed the zebrafish experiments; and R.S.-S. conducted the subcellular localization studies. R.S.S. and D.K.M. wrote the manuscript.

## DECLARATION OF INTERESTS

The authors declare no competing interests.

## STAR METHODS

## LEAD CONTACT AND MATERIALS AVAILABILITY

Further information and requests for resources and reagents should be directed to and will be fulfilled by the Lead Contact, Deborah Morrison. Plasmids and cell lines are available for use upon request to the Lead Contact.

## EXPERIMENTAL MODEL AND SUBJECT DETAILS

### Cell Lines and Culture Conditions

293FT, COS-7, Phoenix-Eco, NIH 3T3, Du1473 and RAS-deficient MEF lines were cultured in DMEM supplemented with 10% fetal bovine serum (FBS), 2 mM L-glutamine, and 1% penicillin/streptomycin. MDCK cells were cultured in EMEM supplemented with 10% FBS. 2 mM L-glutamine, and 1% penicillin/streptomycin. All cells were cultured at 37°C under 5% CO_2_.

### RAF-CRD Protein Production

Wild-type (WT) and mutant human BRAF (232-284) and CRAF/RAF-1 (136-188) entry clones were synthesized with an upstream tobacco etch virus (TEV) protease cleavage site (ENLYFQ/G) and optimized for expression in E. coli by ATUM (Newark, CA). Entry clone inserts were transferred by Gateway recombinational cloning (ThermoFisher, Carlsbad, CA) to pDest-566 (Addgene #11517), a T7-based E. coli expression vector encoding an N-terminal His6-MBP (maltose-binding protein) fusion. All CRD proteins were expressed using LB medium supplemented with 300 mM ZnCl_2_ as outlined for CRAF (52-188) in (Lakshman et al., 2019). For 2-liter working volume fermentations, 3-liter Bioflo 110 (Eppendorf) were used, and 20-liter Bioflo IV (Eppendorf) were used for 15-liter working volumes. Antifoam 204 (Sigma) was added at 0.075% v/v and 0.017% v/v, respectively, to the 2- and 15-liter fermenters. All proteins were purified essentially as outlined in (Kopra et al., 2020) for CRAF RBD (52-131). Specifically, 5 mM TCEP was used in all buffers and the protein occasionally eluted overlapping the salt peak during SEC which then required an additional dialysis step (3K MWCO) to the final buffer.

### SPR measurements

SPR binding experiments were collected on a Biacore T200 Instrument (GE Healthcare). Anti-His6 monoclonal antibody (Abcam) was amine coupled to the carboxy-methylated dextran surface of a CM5 sensor chip (GE Healthcare) using standard amine coupling chemistry. The CM5 chip surface was first activated with 0.1 M N-hydroxysuccinimide and 0.4 M N-ethyl-N’-(3-dimethyl-aminopropyl) carbodimide at a flow rate of 20 µl/min. Next, anti-His6 monoclonal antibody was diluted to 20 µg/ml in 10 mM sodium acetate (pH 5.0) and injected on all 4 flow cells until a density of approximately 10000 Response Units (RU) was attached. His6-tagged MSP1D1 nanodiscs composed of either 1-palmitoyl-2-oleoyl-glycero-3-phosphocholine (POPC) or 80:20 1-palmitoyl-2-oleoyl-glycero-3-phosphocholine:1-palmitoyl-2-oleoyl-glycero-3-phosphoL-serine (POPC:POPS) were prepared as described previously (Agamasu et al., 2019). Nanodiscs were captured (2000-4000 RU) on flow cells 2 and 4 containing the anti-His6 antibody surfaces. Flow cells 1 and 4 were used for referencing and a series of buffer injections were added to establish a stable baseline. BRAF- or CRAF-CRD proteins were diluted in 2-fold from 5 – 0.31 µM in a buffer composed of 20 mM Hepes pH 7.0, 150 mM NaCl and injected over all the flow cells at 30 µl/min. SPR sensorgrams were normalized by the capture level of nanodiscs to allow direct comparison between different experimental runs. Equilibrium binding constants (K_d_) were calculated using Prism. Complementary experiments were performed using captured liposomes as described previously (Travers *et al*., 2018). Specifically, L1 chips were activated with two injections of 20 mM CHAPS followed by liposomes (5 mM) composed of POPC or 80:20 POPC:POPS on flow cells 2 and 4 (capture levels 3500 – 5000). Flow cells 1 and 3 were used for referencing purposes. Following liposome capture a series of buffer injections (20 mM Hepes pH 7.0, 150 mM NaCl) were performed to obtain a stable baseline. BRAF-CRD proteins were diluted 2-fold from 15 – 0.12 µM in buffer and consecutively injected onto the captured liposomes. The L1 was regenerated using two injections 20 mM CHAPS. The data was analyzed to calculate the equilibrium partition coefficient (K_p_; according to (Figueira et al., 2017).

### Transfection and Lysis of Cells

All cell lines were plated 16-20 h prior to transfection at a confluency of ∼70%. Cells were then transfected using the XtremeGENE9 transfection reagent per the manufacturer’s instructions, with a 2:1 ratio of XtremeGENE9 to DNA. For lysis, cells were washed twice with ice cold PBS and lysed in NP40 buffer (20mM Tris [pH 8.0], 137 mM NaCl, 10% glycerol, 1% NP-40 alternative, 0.15 U/mL aprotinin, 1 mM phenylmethylsulfonyl fluoride, 0.5 mM sodium vanadate, 20 mM leupeptin) at 4°C for 15 min on a rocking platform. Lysates were clarified by centrifugation at 14,000 rpm for 10 min at 4°C, following which the protein content was determined by Bradford assays.

### Cell Imaging

COS7 and MDCK cells (5x10^4^) were plated onto 35 mm glass-bottom dishes (MATEK) that had been pre-coated with 10 µg/ml human placental type IV collagen. 16 h after plating, cells were transfected with the indicated DNA constructs, incubated for 24 h at 37°C under 5% CO_2_, and then serum-starved overnight, where indicated. The following day, cells were washed twice with PBS, fixed for 10 min in 4% paraformaldehyde, and then washed twice with PBS. MDCK cells were permeabilized with 0.1% Triton-PBS for 15 min, washed twice with PBS, blocked for 1 h in 2% BSA-PBS, and then incubated overnight in a 0.1% BSA-PBS solution containing E-cadherin antibody. The following day, cells were washed three times with PBS, incubated with Alexafluor568 secondary antibody for 45 min, washed three times with PBS containing 0.2% Tween-20 (TBST), and then washed twice in PBS. and then imaged in PBS using a 63x oil objective on a Zeiss LSM710 microscope. For the quantification of plasma membrane enrichment, cells were selected at random in the red (E-cadherin) channel, linear profiles were drawn through the plasma membrane and into the cytosol using ImageJ, the fluorescent intensity of both the red and green (BRAF^FL^-Venus) channels was measured, and the background fluorescence subtracted. Percent plasma membrane enrichment of the BRAF^FL^-Venus was calculated by dividing the peak plasma membrane fluorescence by the average of a 2 µm cytosolic section in the green channel (BRAF^FL^-Venus) for a total of 20 cells per condition.

### NanoBRET Autoinhibition Assay

293FT cells were seeded into 6-well tissue culture plates at a concentration of 4x10^5^ cells/well. 16 h after plating, cells were co-transfected using 5 ng of the indicated NanoLuc-RAF^CAT^ construct and 20 ng of the indicated Halo-BRAF^REG^ construct (per well). 24 h later, cells were collected and resuspended in serum-free/phenol free Opti-MEM (Gibco^TM^). 1 µl/ml HaloTag NanoBRET^TM^ 618 ligand was added to the cell suspension and cells were seeded at a concentration of 8 x10^3^ cells per well of a 384-well plate in quadruplicate (BioTek, Winooski, VT), with the remaining cells plated into a fresh 6-well tissue culture plate. After 24 h, 10 µl/ml NanoBRET NanoGlo substrate was added to each well of cells to be monitored for NanoBRET, with donor (460nm) and acceptor (618nm) emissions measured using a Perkins Elmer Envision plate reader (#2104-0010A containing a 460nm/50nm emission filter and a 610nm LP filter). Cells seeded into the 6-well plates were lysed, and the lysates examined by immunoblot analysis using antibodies recognizing the Halo-tag (for RAF^REG^ detection) or the NanoLuc-tag (for RAF^CAT^ detection) to ensure equal expression levels across conditions.

### BRET RAS/RAF Interaction Assay

293FT cells were seeded into 12-well dishes at a concentration of 1x10^5^ cells/well. 16 h after plating, Venus-tagged and Rluc8-tagged constructs were co-transfected into cells using a calcium phosphate protocol. Duplicate 12-point saturation curve was generated in which the concentration of the energy donor construct (Rluc8) was held constant (62.5 ng) as the concentration of the energy acceptor plasmid (Venus) increased (0-1.0 μg). Cells were collected 48 h after transfection, washed, and resuspended in PBS (500 μL). 30 μL of the cell suspension was plated in duplicate into wells of a 384-well white-walled plate (PerkinElmer CulturPlate) and coelenterazine-h was added to a final concentration of 3.375 μM. The BRET and Rluc8 emissions were measured simultaneously using a PHERAstar Plus plate reader (BMG Labtech), with BRET monitored at 535 nm (bandwidth 30 nm) and Rluc8 measured at 475 nm (bandwidth 30 nm). To monitor the increasing levels of acceptor expression, 90 μL of the cell suspension was also plated in duplicate into wells of a 96-well black-walled plate (PerkinElmer OptiPlate), and Venus fluorescence was determined by using an excitation wavelength of 485 nm (5 nm bandwidth) and monitoring the emission at 530 nm (5 nm bandwidth) on a Tecan Infinite M1000 plate reader. The BRET ratio was determined by first calculating the 535/475 ratio for a given sample (sample BRET) and then normalizing the sample BRET to the BRET signal obtained from cells expressing the donor construct alone. The normalized mBRET signal was calculated as follows: (1000 X (sample BRET ratio/background BRET ratio)) – 1000. The acceptor/donor ratio (Venus emission/Rluc8 emission) for each data point was background corrected and equalized against a control where equal quantities of the Venus (acceptor) and Rluc8 (donor) constructs were transfected. Data was analyzed using GraphPad Prism. Non-linear regression was used to plot the best fit hyperbolic curve, and values for BRET_max_ and BRET_50_ were obtained from the calculated best fit curves.

### Focus Forming Assay

Recombinant retroviruses expressing the RAF proteins of interest were generated by transfecting 6 μg of the desired DNA construct into Phoenix-Eco cells (100 mm dish) using the XtremeGENE9 protocol described above. Viral supernatants were collected 48 h post-transfection, centrifuged twice at 1500 rpm for 10 min, and either stored at -80°C or used directly. NIH-3T3 cells were plated into 60 mm tissue culture dishes at a concentration of 2 x 10^5^ cells/dish. After 16 h, cells were infected with the indicated recombinant retrovirus in media containing 4% FBS and 8 μg/mL polybrene. 24 h later, cells were trypsinized and plated into two 100 mm dishes, one of which contained 5 μg/mL puromycin. After 2-4 weeks of culture, cells were fixed with 3.7% formaldehyde and stained with 1% methylene blue.

### Proliferation Assay

Recombinant retrovirus expressing the indicated BRAF^FL^ variants were generated as described for focus forming assays. HRAS and NRAS-deficient MEFs (DU1473) were infected or not with the recombinant retroviruses, following which cells were selected for 7 days in media containing 2 ug/ml puromycin. Resistant lines were then treated for 14 days with 500 nM doxycycline to remove the floxed KRAS-allele. Lysates from the doxycycline-treated cells were then examined by immunoblot analysis using the pan-RAS antibody (ABCAM) to confirm the absence of RAS protein expression. Cells were plated at a concentration of 1x10^3^ cells/well into 24-well tissue culture plates and proliferation was assessed every 24 hours over a 4-day time course using CellTiter-Glo 2.0 reagent (Promega). Luminescence was measured using a Perkins Elmer Envision Multimode Plate Reader.

### Zebrafish Embryo Injections and Analysis

Fish care and animal work was approved by the National Cancer Institute at Frederick Animal Care and Use committee (Study Proposal 17-416). All capped mRNAs were generated from NotI linearized pCS2+ vector templates using the mMESSAGE mMACHINE T3 kit (ThermoFisher Scientific). AB-Tübingen zebrafish embryos were microinjected at the one cell stage with 66 pg/nL of mRNA encoding either WT or mutant FLAG-BRAF proteins or WT or mutant BRAF-Venus fusion proteins. After injection, embryos were maintained at 25°C until analysis. For convergence-extension cell movement assays (CE assays), embryos at the 3-6 somite stage were embedded oriented laterally in 1% low-melt agarose (Invitrogen, Carlsbad, CA) and imaged under bright field (to determine yolk elongation by taking major and minor axis measurements) and under fluorescent conditions (to monitor expression of the BRAF-Venus proteins). Larvae collected three days post-fertilization (dpf) were positioned laterally and imaged under bright field to monitor for pericardial edema. For ceratohyal (CH) angle and length measurements, Tg (*col2a1aBAC:GFP*) embryos were injected with 33 pg/nL RNA and treated with 0.003% 1-phenyl 2-thiourea to block pigmentation. At 5dpf, larvae were anesthetized with 0.2 mg/mL Tricaine in E3 and positioned dorsally against the side of grooves made by a mold (World Precision Instruments) in 1% low-melt agarose.

Images under bright field condition were taken using a 2X water immersion objective (0.06 NA) for CE assays, and image acquisition (∼ 200 mm z-stacks with 5 mm step size) of Venus fluorescence was performed using a 10X objective (0.3 NA) for CH analyses. The Nikon Eclipse Ni-E upright microscope was equipped with a DS-Ri2 camera. All measurements were performed using the Nikon NIS Elements Analysis software and processing of images were carried out using ImageJ and Photoshop. Statistical analyses were conducted using GraphPad Prism.

### RAF Kinase Assay

Cells expressing the indicated FLAG-RAF proteins were lysed under stringent conditions in RIPA buffer (20mM Tris [pH 8.0], 137 mM NaCl, 10% glycerol, 1% NP-40 alternative, 1% SDS, and 0.1% sodium deoxycholate) to disrupt protein complexes. The RAF proteins were immunopurified and incubated in kinase buffer (30 mM Tris pH 7.4, 10 mM MnCl2, 5 mM MgCl2, 1 mM DTT, 16.5 μM ATP) containing 20 μCi [γ-32P]ATP and kinase-inactive MEK. Samples were resolved by SDS-PAGE, and after transfer to nitrocellulose, the amount of radioactivity incorporated into MEK was determined by scintillation counting. RAF activity levels were normalized to RAF protein levels.

## KEY RESOURCES TABLE

| REAGENT or RESOURCE | SOURCE | IDENTIFIER |
| --- | --- | --- |
| **Antibodies** |  |  |
| BRAF (H-145) rabbit polyclonal | Santa Cruz Biotechnology | cat# sc-9002; RRID:AB_2067494 |
| HaloTag mouse monoclonal | Promega | cat# G9211; RRID:AB_2688011 |
| NanoLuc (Nluc) luciferase antibody | R&D Systems | cat# MAB100261 |
| FLAG M2 antibody | Millipore/Sigma | cat# F3165; RRID:AB_439685 |
| FLAG (L5) rat monoclonal antibody | BioLegend | cat# 637304; RRID:AB_1134267 |
| MEK1 mouse monoclonal | BD Biosciences | cat# 61022; RRID:AB_397528 |
| pS217/221-MEK rabbit polyclonal | Cell Signaling Technology | cat# 9121; RRID:AB_331648 |
| β-Tubulin mouse monoclonal | Santa Cruz Biotechnology | cat# sc-5274; RRID:AB_2288090 |
| α-Tubulin antibody, mouse monoclonal | Millipore Sigma | cat# T6199; RRID:AB_477583 |
| Pan-RAS rabbit polyclonal antibody | Cell Signaling Technology | cat# 3965; RRID:AB_2180216 |
| Alexa Fluor 568 goat anti-rat polyclonal antibody | Thermo Fisher | cat# A-11077; RRID:AB_2534121 |
| E-Cadherin, rat monoclonal antibody | Invitrogen | cat# 14-3249-82; RRID:AB_1210458 |
| **Chemicals** |  |  |
| Platelet Derived Growth Factor (PDGF) | Millipore/Sigma | cat# SRP3229 |
| X-tremeGENE 9 | Roche/Sigma | cat# 06365809001 |
| Collagen, Human Placenta Type IV | Millipore/Sigma | cat# C7521 |
| Doxycycline ((Z)-4-Hydroxytamoxifen) | Millipore Sigma | cat# H7904 |
| Paraformaldehyde | EM Sciences | cat# 15710 |
| Coelenterazine-h | Promega | cat# S2011 |
| **Critical Commercial Assays** |  |  |
| CellTiter-Glo 2.0 | Promega | cat# G9241 |
| QuickChange II Kit Agilent | Agilent | cat# 200523 |
| NanoBRET Kit | Promega | cat# N1661 |
| **Experimental Models: Cell Lines** |  |  |
| 293FT (human) | ATCC | cat# PTA-5077; RRID:CVCL_6911 |
| Phoenix-Eco (human) | ATCC | cat# CRL-3214; RRID:CVCL_H717 |
| NIH 3T3 (mouse) | ATCC | cat# CRL-1658; RRID:CVCL_0594 |
| COS7 | ATCC | cat# CRL-1651; RRID:CVCL_0224 |
| DU1473 | NCI-RAS Initiative | N/A |
| **Recombinant DNA** |  |  |
| pEGFP-C1-BRAF-CRD mutants | This paper | N/A |
| pCMV5-BRAF<sup>REG</sup>-mutants-Halo | This paper | N/A |
| pCMV5-NanoLuc-BRAF<sup>CAT</sup> mutants | This paper | N/A |
| pCMV5-NanoLuc-CRAF<sup>CAT</sup> | This paper | N/A |
| pCMV5-CRAF<sup>REG</sup>-Halo | This paper | N/A |
| pLenti-KRAS<sup>G12V</sup> | NCI-RAS Initiative | N/A |
| pCMV5-Venus-KRAS<sup>G12V</sup> | NCI-RAS Initiative | N/A |
| pLHCX-BRAF<sup>FL</sup>-RLuc8 | Terrell *et al.*, 2019 | N/A |
| pLHCX-BRAF<sup>FL</sup> CRD mutants-RLuc8 | This paper | N/A |
| pLHCX-BRAF<sup>REG</sup> CRD-RLuc8 | Terrell *et al.*, 2019 | N/A |
| pLHCX-BRAF<sup>REG</sup>CRD mutants-RLuc8 | This paper | N/A |
| pCS2+-BRAF<sup>FL</sup> CRD mutants | This paper | N/A |
| pBabe-Puro-FLAG-BRAF | Freeman *et al.*, 2012 | N/A |
| pBabe-Puro-FLAG-BRAF mutants | This paper | N/A |
| pBabe-Puro-FLAG-CRAF | Freeman *et al.*, 2012 | N/A |
| pBabe-Puro-FLAG-CRAF mutants | This paper | N/A |
| pEGFP-C1-CRAF-CRD mutants | This paper | N/A |
| **Software and Algorithms** |  |  |
| GraphPad Prism | GraphPad | https://www.graphpad.com |
| ImageJ | ImageJ | https://imagej.nih.gov/ij/ |

## SUPPLEMENTAL FIGURES AND TABLES

**Figure S1, related to Figure 1.**
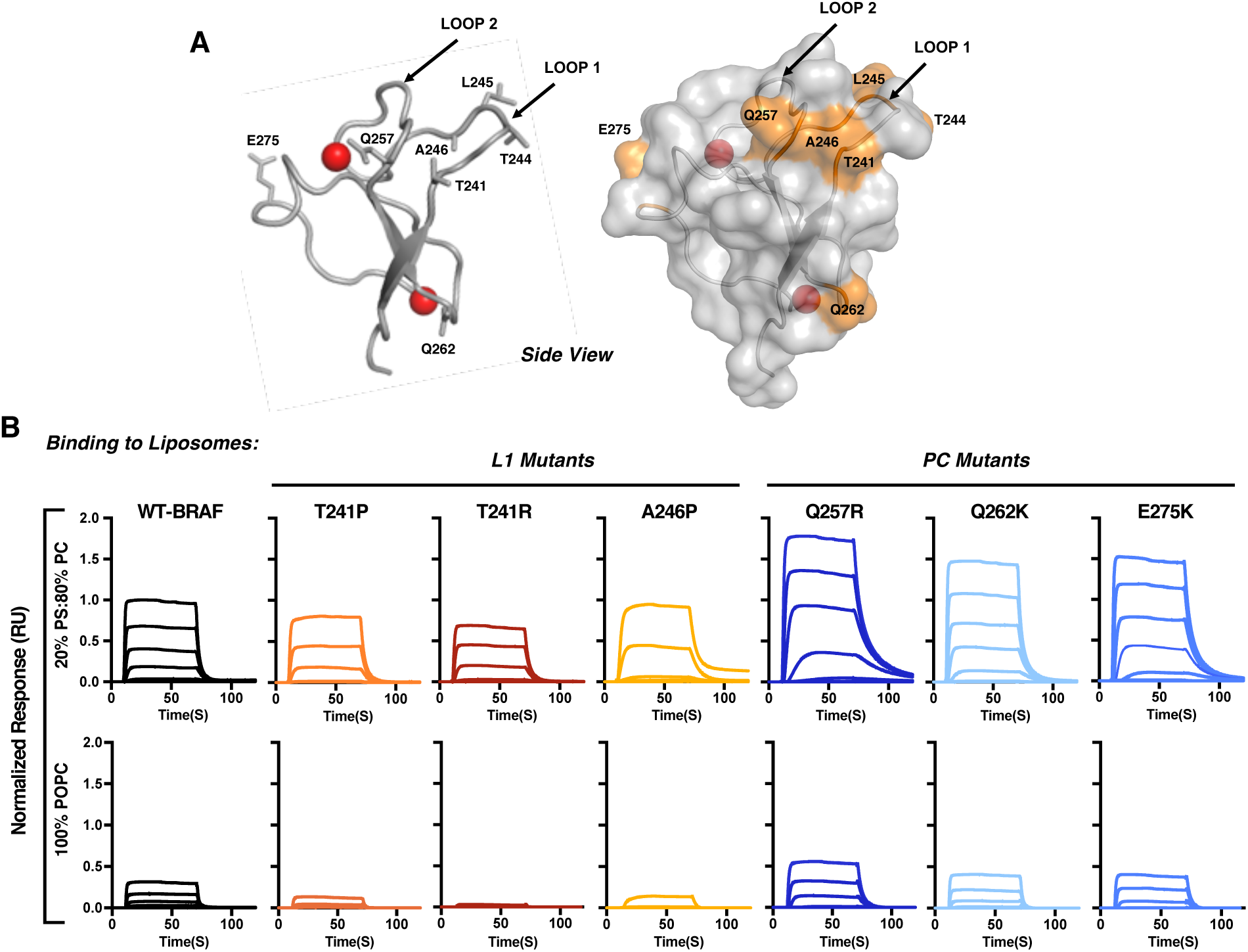
Analysis of the Lipid Binding Properties of the BRAF-CRD Mutants. (A) Ribbon (left) and space filling (right) diagrams of the BRAF-CRD with the coordinated two Zn^2+^ ions depicted as red spheres. CRD residues mutated in the RASopathies are labeled and shown in orange in the space filling diagram and their side chains are shown in the ribbon diagram (PDB: 6NYB). (B) A 2-fold dilution series of purified 5 µM WT or mutant BRAF-CRD proteins was injected over liposomes containing 20% POPS/80% POPC or 100% POPC, and binding responses were determined by SPR. Sensorgrams were normalized to the WT-BRAF-CRD binding response. Binding responses are representative of 3 independent experiments with similar results.

**Figure S2, related to Figure 2.**
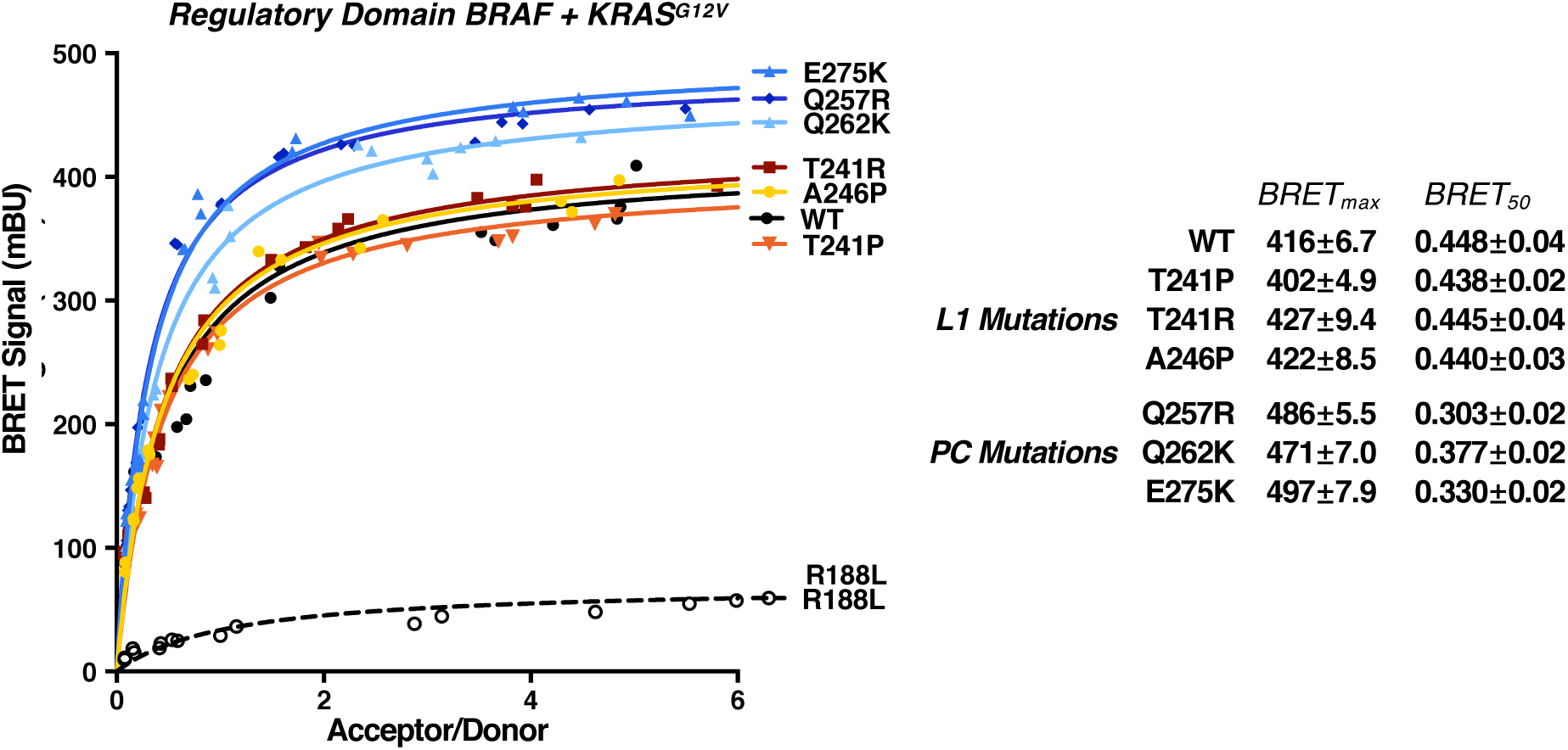
Effect of BRAF-CRD Mutations on Binding of BRAF^REG^ to KRAS^G12V^ in Live Cells. BRET saturation curves are shown examining the interaction of WT or mutant BRAF^REG^-RLuc8 proteins with Venus-KRAS^G12V^. BRET_max_ and BRET_50_ values derived from the saturation curves are also listed ± SEM. Donor and acceptor expression levels were monitored via the RLuc8 and Venus emissions and are incorporated into the BRET calculations. Saturation curves were repeated 3 times with similar results.

**Figure S3, related to Figure 3.**
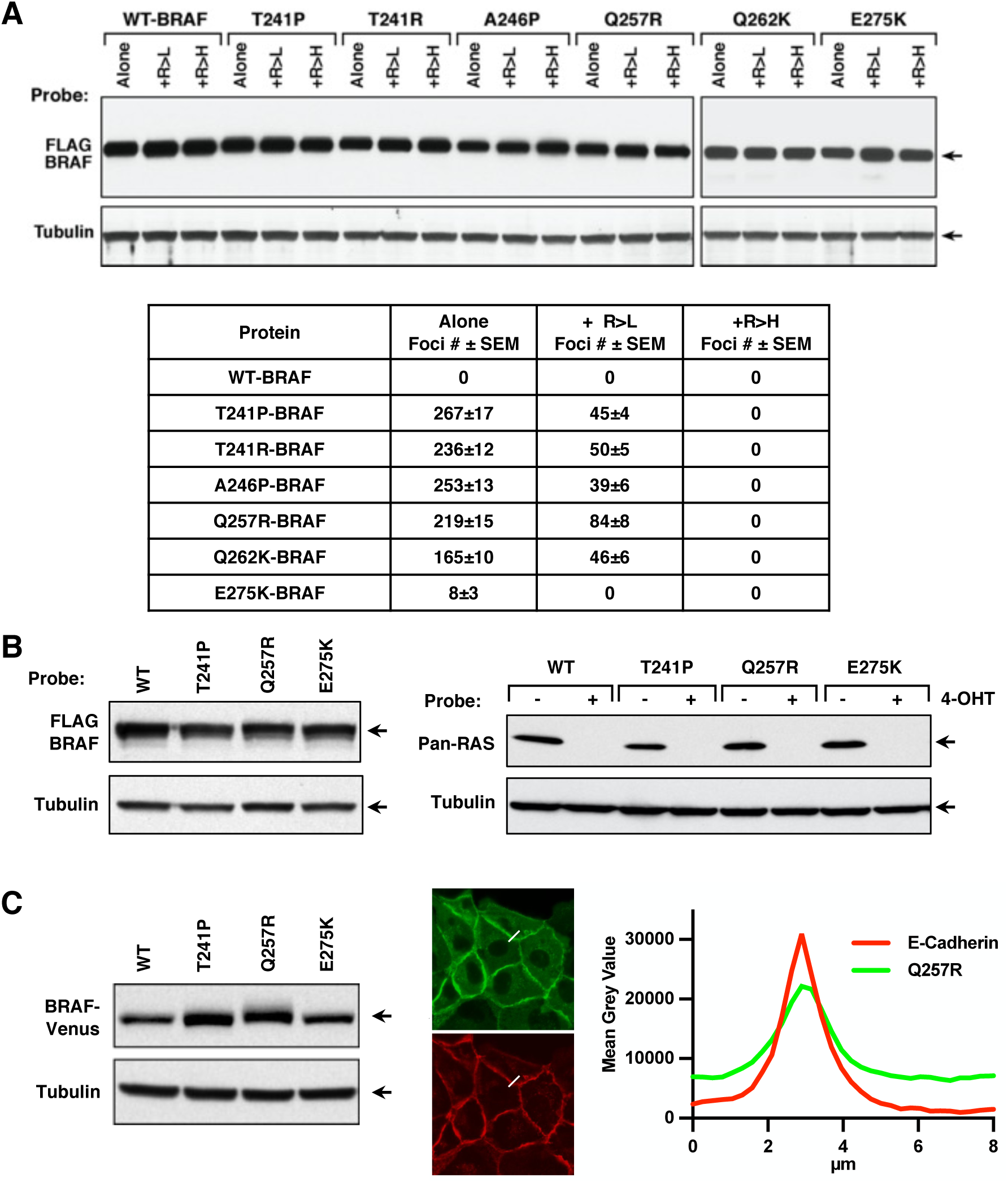
Analysis of BRAF-CRD Mutants in NIH-3T3 Cells, RAS-deficient MEFs, and MDCK Cell Lines. (A) Representative plates in the NIH-3T3 focus assays were taken for cell lysis, and lysates were examined by immunoblot analysis for expression of the indicated FLAG-BRAF proteins and tubulin (upper). Quantification of the foci number ± SEM from 3 independent assays is also shown (below). (B) DU1473 cell lysates were examined by immunoblot analysis to demonstrate the equivalent expression of the FLAG-BRAF^FL^ variants (left) and the lack of endogenous RAS proteins (right). (C) MDCK cell lysates were examined by immunoblot analysis to demonstrate the equivalent expression of the indicated BRAF^FL^-Venus proteins (left panel). Example confocal images of serum-starved MDCK cells expressing Q257R-BRAF-Venus that were used to quantify the plasma membrane recruitment of BRAF. Also, shown is a tracing depicting the fluorescence intensity of BRAF-Venus (green) and E-cadherin (red) in the area indicated by the white line (right panel).

**Figure S4, related to Figure 5.**
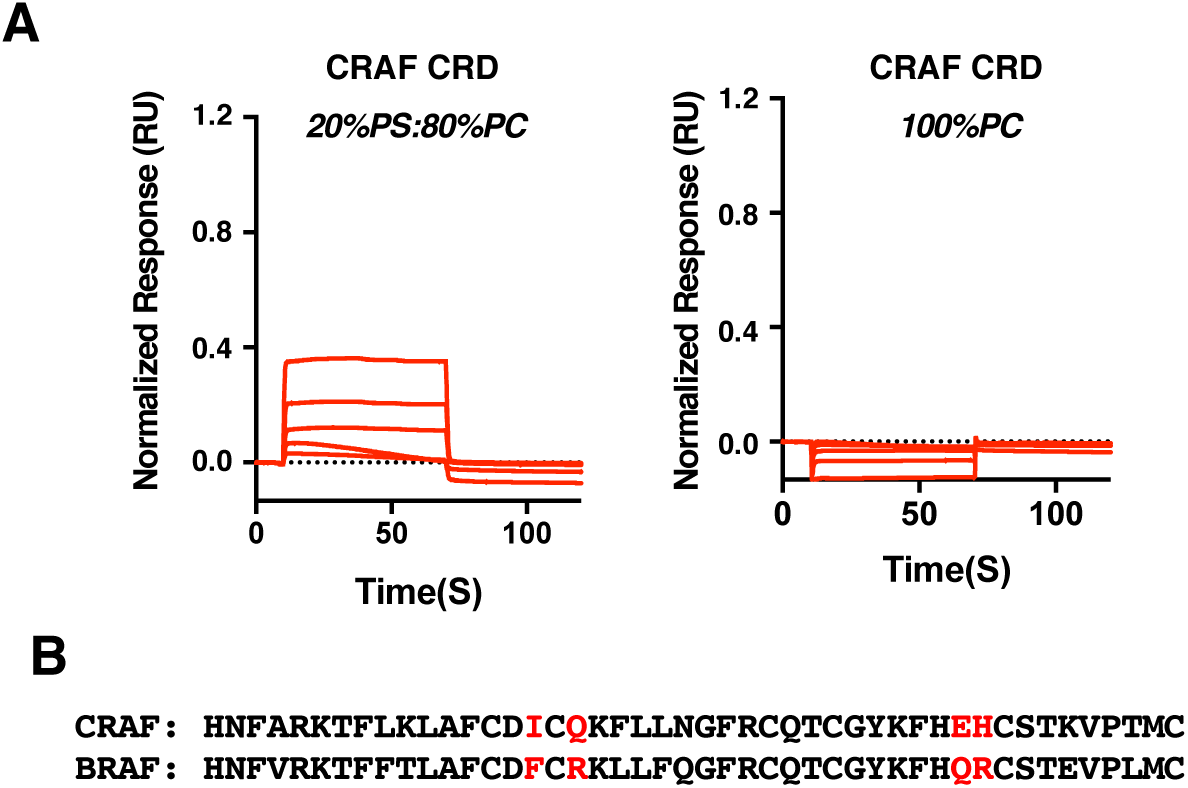
The CRAF-CRD Exhibits Preferential Binding to PS-containing Nanodiscs. (A) A 2-fold dilution series of purified 5mM CRAF-CRD was injected over nanodiscs containing 20% POPS/80% POPC or 100% POPC, and binding responses were determined by SPR. (B) Shown is the sequence alignment of the BRAF and CRAF-CRDs with the 4MUT residues shown in red.

**Table S1, related to Figures 1-4.** Summary and Classification of BRAF-CRD Mutants.

|  |  | *L1 Mutations* |  |  | *PC Mutations* |  |  |
| --- | --- | --- | --- | --- | --- | --- | --- |
|  | WT | T241P | T241R | A246P | Q257R | Q262K | E275K |
| Lipid Binding | Yes | ~ to WT | ~ to WT | ~ to WT | ↑↑↑ | ↑↑ | ↑↑ |
| Autoinhibition | Yes | ↓↓ | ↓↓ | ↓↓ | ↓↓ | ↓↓ | ~ to WT |
| CLASS |  | A | A | A | B | B | C |
| KRAS<sup>G12V</sup> binding<br>FL-BRAF | Yes | ↑↑ Levels<br>↑ Affinity | ↑↑ Levels<br>↑ Affinity | ↑↑ Levels<br>↑ Affinity | ↑↑ Levels<br>↑↑ Affinity | ↑↑ Levels<br>↑ Affinity | ↑ Levels<br>↑ Affinity |
| PM Localization<br>FL-BRAF | Primarily<br>Cytosolic | ↑ Starved<br>↑↑ Cycling | nd | nd | ↑↑ Starved<br>↑↑↑ Cycling | nd | ~WT Starved<br>↑ Cycling |
| Transformation | No | Yes | Yes | Yes | Yes | Yes | Yes, Weaker |
| Dimer-dependent | Yes | Yes | Yes | Yes | Yes | Yes | Yes |
| RAS-dependent | Yes | ↓ Reduced | ↓ Reduced | ↓ Reduced | ↓↓ Reduced | ↓ Reduced | Yes |
| Zebrafish defects | No | Yes | nd | nd | Yes | nd | Yes, Weaker |

*Class A – mutation relieves autoinhibition*
*Class B – mutation relieves autoinhibition and enhances PS/membrane binding*
*Class C – mutation enhances PS/membrane binding but has no effect on autoinhibition*

**Table S2, related to Figure 4.** Reported Evaluations of Patients with BRAF-CRD Mutations. See attached Excel file.

**Table S3, related to Figure 6.** Effect of CRD Residue Swaps on BRAF Autoinhibitory Interactions.

| BRAF<sup>Reg</sup>-Halo Construct | Reduced Autoinhibition |
| --- | --- |
| **CRD swap** | +++ |
| **T244K** | +++ |
| **V238A/T244K** | +++ |
| **F243L/T244K** | +++ |
| **V238A/F243L/T244K** | ++++ |
| **F250I** | ~15% increase |
| **R252Q** | No |
| **F250I/R252Q** | No |
| **F250I/R252Q/L254F/F256L/Q257N** | +++ |
| **F250I/R252Q/Q270E/R271H** | No |
| **E275K** | No |
| **Q270E/R271H/E275K** | No |
| **Q270E/R271H/E275K/L278T** | ++ |
| **T244K/R252Q** | +++ |
| **T244K/L254F** | +++ |
| **T244K/F256L** | ++++ |
| **T244K/Q257N** | +++ |

## Notes

### Competing Interest Statement

The authors have declared no competing interest.

### Summary of Updates

This is a revised version of the manuscript, which includes changes to the title, abstract, main text and figures.

